# Transposable elements facilitate the unintentional domestication of a cheese-associated *Penicillium* mold

**DOI:** 10.1101/2025.04.10.648237

**Authors:** Nicolas L. Louw, Justin L. Eagan, Jackson Larlee, Mateo Kehler, Nancy P. Keller, Benjamin E. Wolfe

**Affiliations:** Tufts University, Department of Biology, Medford, MA, 02155; University of Wisconsin-Madison, Medical Microbiology & Immunology, Madison, WI, 53706; Jasper Hill Farm, Greensboro Bend, VT, 05842; University of Wisconsin-Madison, Department of Plant Pathology, Madison, WI, 53706

**Author notes:** Author for correspondence: Benjamin E. Wolfe.

**Keywords:** *Penicillium*, fungi, adaptation, melanin, relaxed selection, cheese, transposable elements

## Abstract

Previous comparative and experimental evolution studies have suggested how fungi may rapidly adapt to new environments, but direct observation of *in situ* selection in fungal populations is rare due to challenges with tracking populations over human time scales. We monitored a population of *Penicillium solitum* over eight years in a cheese cave and documented a phenotypic shift from predominantly green to white strains. Diverse mutations in the *alb1* gene, which encodes the first protein in the DHN-melanin biosynthesis pathway, explained the green to white shift. A similar phenotypic shift was recapitulated with an *alb1* knockout and experimental evolution in laboratory populations. The most common genetic disruption of the *alb1* genomic region was caused by putative transposable element insertions upstream of the gene. White strains had substantial downregulation in global transcription, with genetically distinct white strains possessing divergent shifts in expression of different biological processes. White strains outcompeted green strains in co-culture, but this competitive advantage was only observed in the absence of light, suggesting that loss of melanin is only adaptive in dark conditions. Our results illustrate how fermented food production by humans provides opportunities for relaxed selection of key fungal traits over short time scales. Unintentional domestication of microbes by cheesemakers may provide opportunities to generate new strains for innovation in traditional cheese production.

## INTRODUCTION

Understanding how fungi adapt to different environments can help us better predict pathogen evolution, develop strains for biotechnology, and identify how fungal populations might respond to environmental changes ^1–5^. Most studies of fungal adaptation use comparative genomic approaches that rely on extant diversity and past events to identify mechanisms of fungal adaptation ^6–12^. These are powerful approaches that can infer potential evolutionary patterns and processes in fungi, but cannot directly reveal the mechanisms by which fungal traits evolve in populations over short time scales ^13^. Experimental evolution studies can directly reveal rates and modes of fungal trait evolution ^14–17^, but the population dynamics and laboratory environments used in these studies do not necessarily reflect those in natural environments ^18^.

Cheeses that are aged in caves provide natural colonization experiments to observe selection within fungal populations over human time scales. When surface-ripened cheeses are placed in natural or artificial cave environments, they form microbial rinds on their surfaces composed of communities of bacteria, yeasts, and filamentous fungi (molds) ^19,20^. Microbial populations from soil, plant, and marine environments can colonize and adapt to the environments of cheese caves ^21–23^. Past comparative genomic and experimental evolution studies have provided various lines of evidence for potential domestication by humans of multiple cheese-associated *Penicillium* molds ^7,24^, but how quickly and through what mechanisms fungi adapt to cheese in actual aging environments has not been directly identified.

In this work, we leveraged our long-term collaboration with an artisan cheesemaker in Vermont (USA)^25^ to track the *in situ* evolution of a cheese-associated fungal population (*Penicillium solitum*). Using phenotypic assays, metagenomics, comparative genomics, fungal genetics, transcriptomics, experimental evolution, fitness assays, and microbiome reconstructions, we provide a comprehensive portrait of how cheese-associated fungal populations can adapt to cheese cave environments by strongly shifting their biology. Unlike comparative approaches that can only infer how domestication *may* have unfolded, we were able to directly watch and carefully document the unintentional domestication of a mold over a decade of cheese production.

## RESULTS

### A cheese cave population of *Penicillium solitum* shifted from green to white phenotypes over a decade

At the Cellars at Jasper Hill cheese aging facility in Vermont, we noticed a dramatic shift in the appearance of the rind of a naturally-aged blue cheese called Bayley Hazen Blue (**Figure 1A**). This cheese is inoculated with an interior mold (*P. roqueforti*) to develop veins and is placed on wooden shelves in a dark, cool, humid cave made from concrete. Over a period of about 60 days, it develops a natural rind microbial community on the surface of the cheese ^19,20^. Natural rinds can impart flavors during aging and can affect cheese aesthetics by forming a distinct coating on the external surface of the cheese ^26,27^. Microbes that form the rind, including bacteria, yeasts, and molds such as *Penicillium* species, can come from raw milk and other raw materials (e.g. salt), starter cultures, and the cheese cave environment ^21–23^.

**Figure 1:**
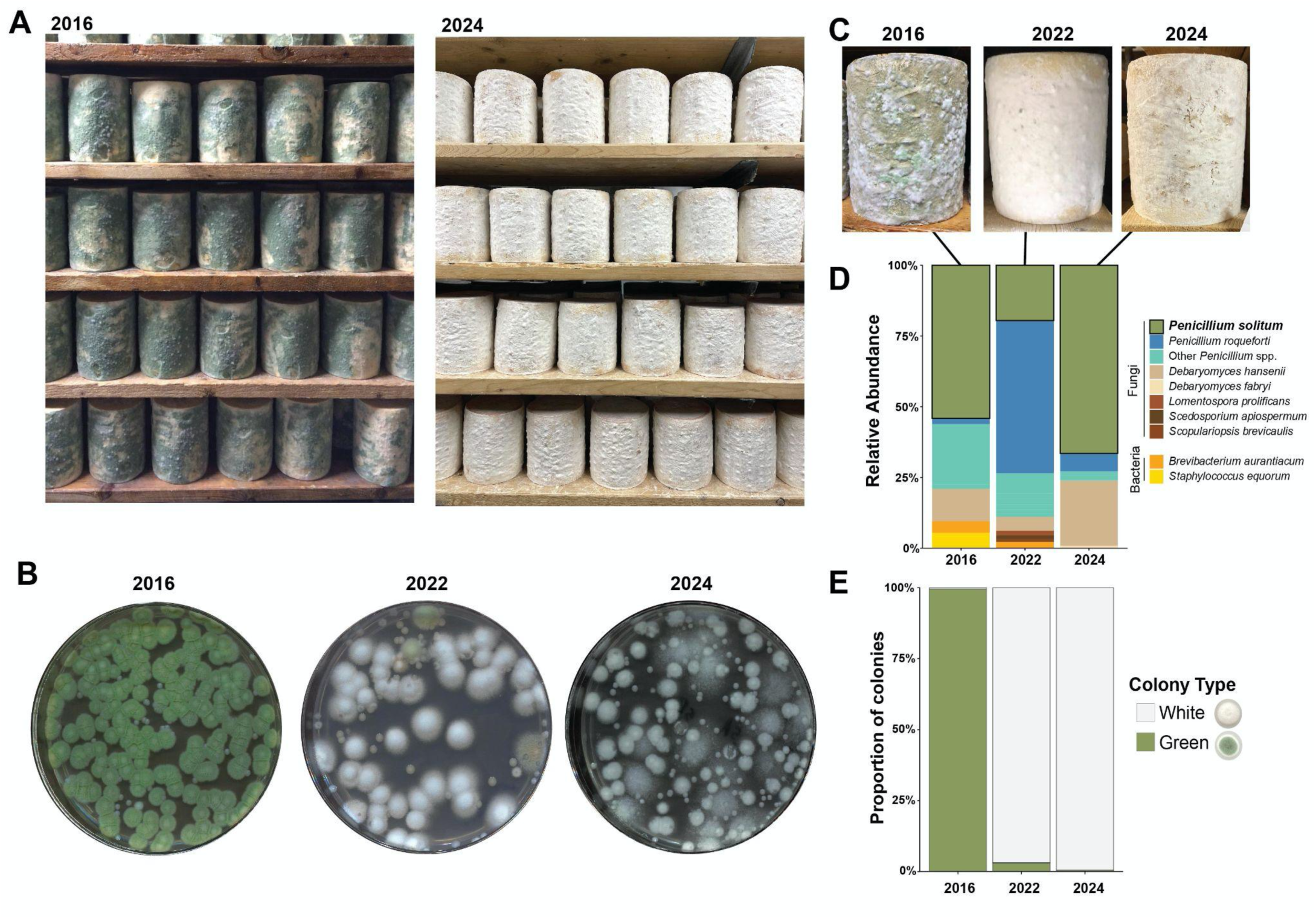
A white phenotype displaces a green phenotype of *Penicillium solitum* in a cheese aging facility. **(A)** Photographs taken at the same cheese aging vault in 2016 (left) and 2024 (right) portraying a phenotypic shift across eight years. See Figure S1 for additional photos and time points. **(B)** Representative images of fungal colonies from a cheese rind sample from each of the different years sampled. Plates contain plate count agar with milk and salt (PCAMS). Medium-sized white colonies in 2022 and 2024 are white phenotypes of *P. solitum*. Small white colonies are the yeast *Debaryomyces hansenii* and large colonies in 2024 are a *Scopulariopsis* species. **(C)** Photographs of representative wheels of cheese from each year. **(D)** Taxonomic assignment from the metagenomic classifier kaiju of the most abundant microbial taxa detected with shotgun metagenomic sequencing. Each stacked relative abundance bar represents the mean of three samples taken from separate wheels in each year. **(E)** The proportion of white to green *P. solitum* phenotypes across the sampling years. Each stacked relative abundance bar represents the proportion of white to green phenotypes of *P. solitum* within each sampling year, calculated as the total number of white and green colonies counted within each year across all 50 samples taken in each year.

In our early years of studying this cheese (2012-2016), the rind had a mint green appearance and was dominated by a mix of filamentous fungi (**Figure 1A**). In our previous work, we noted that this green rind was formed by the fungus *P. solitum*, and that there was a very low frequency of white strains within the population ^28^. We do not know the original source of this *P. solitum* population; this species is sometimes detected on cheese ^29^, but it is not sold as a starter culture and is commonly found associated with apples and marine environments ^30,31^.

At some point between 2016 and 2022 (**Figure S1**), there was a striking shift in the appearance of the rind from green to white, and the white phenotype persists to the time of this publication (**Figure 1A-C**). We hypothesized that this shift could be caused by either extreme changes in the species composition of the rind community, or an alteration in the phenotype of a dominant fungus in the rind. To distinguish between these two potential causes, we used a set of samples collected in 2016, 2022, and 2024 (one rind sample from each of 50 distinct wheels of cheese throughout the cave) to characterize the species composition of the rind community using culture-dependent (plating to observe morphologies of dominant molds) and independent (shotgun metagenomics) approaches. These samples were not specifically collected for a temporal study of fungal adaptation, but were part of other cheese microbial ecology projects in the lab. These were frozen as glycerol stocks and provided a microbial time capsule to understand community and phenotypic changes in this rind community.

Culturing identified *P. solitum* as the dominant fungus in 100, 73, and 100% of samples plated out from 2016, 2022, and 2024, respectively (**Figure 1B**). Beyond *P. solitum*, other fungal taxa included various *Penicillium* species, *Scopulariopsis* species, and the yeast *Debaryomyces hansenii*, which are typical members of cheese rind microbiomes ^25^. To further confirm that *P. solitum* was a dominant community member, we selected three samples per year for shotgun metagenomic sequencing to measure the abundance of different microbial taxa in the rinds (**Table S1**). Of the classified reads, *P. solitum* was the most abundant non-starter (excluding the blue mold *P. roqueforti* that is added to the cheese) fungal species across the three different years (**Figure 1D; Figure S2**). These plating and metagenomic data indicate that a shift in fungal community composition does not explain the rind phenotype shift from green to white.

When quantifying the phenotypes of *P. solitum* colonies across the three different sampling times, we noticed a clear pattern that aligned with shifts in the cheese appearance; the proportion of white colonies compared to green changed from <1% (2016), to 97% (2022) and 99.6% (2024), signifying an almost complete phenotypic shift from green to white during the sampling period (**Figure 1E**). The sampling scheme was not originally designed for temporal sampling, and we acknowledge that we have a large temporal gap in our sampling between 2016 and 2022. However, additional photographic documentation of the rinds from the cheese producers and social media between 2016 and 2022 indicates a mix of green and white appearance in 2018 and 2019 (**Figure S1**). Without samples taken at these time points, we are unable to identify the exact rate of phenotypic changes in this population. But our direct sampling from three years and our additional photographic evidence suggests that there has been a substantial and lasting phenotypic shift in this *P. solitum* population.

### White *Penicillium solitum* strains have mutations in the DHN-melanin pathway

To identify potential genetic mechanisms causing the striking shifts in *P. solitum* phenotypes over time, we sequenced genomes of 43 strains (11 green and 32 white) isolated from across the three sampling times (**Table S2**). We used a combination of Oxford Nanopore long-read and Illumina short-read sequencing to create a contiguous and high quality reference genome of the *P. solitum* ancestor. For the reference genome, we used a green *P. solitum* strain that was isolated in 2012 from the same location and has been extensively characterized in previous work ^25,32,33^. We refer to this isolate throughout as the “ancestor” *P. solitum* or “*P. solitum* strain #12” in downstream analyses and experiments. We acknowledge that this single isolate does not fully capture the genetic diversity of the initial colonizing population of *P. solitum*, but it provided a useful reference for comparing all 2016, 2022, and 2024 genomes. We used short-read re-sequencing (Illumina) on *P. solitum* isolates collected over the three years to identify single nucleotide polymorphisms (SNPs) and short indels in coding regions that might help explain the green to white phenotypic shift.

The number of non-synonymous mutations ranged from three to five mutations per green and white strain, with significantly more SNPs on average in white strains (F_1,40_ = 4.9, *p* = 0.03; **Figure S3**). Using an ordination of all 97 high-confidence SNPs observed across all strains, we observed that most of the 2016 green strains clustered together (**Figure 2A; Table S3**). The only white strain to cluster with these green strains (35A) is a white isolate that was experimentally evolved from a green strain (see below). Interestingly, all green strains isolated in 2022 (82_G, 85_G) and one green strain isolated in 2016 (1_G) clustered with many white strains (**Figure 2A**). Based on their many shared SNPs (**Table S3**), these green strains may be the precursors of many of the white strains isolated in 2022 and 2024.

**Figure 2:**
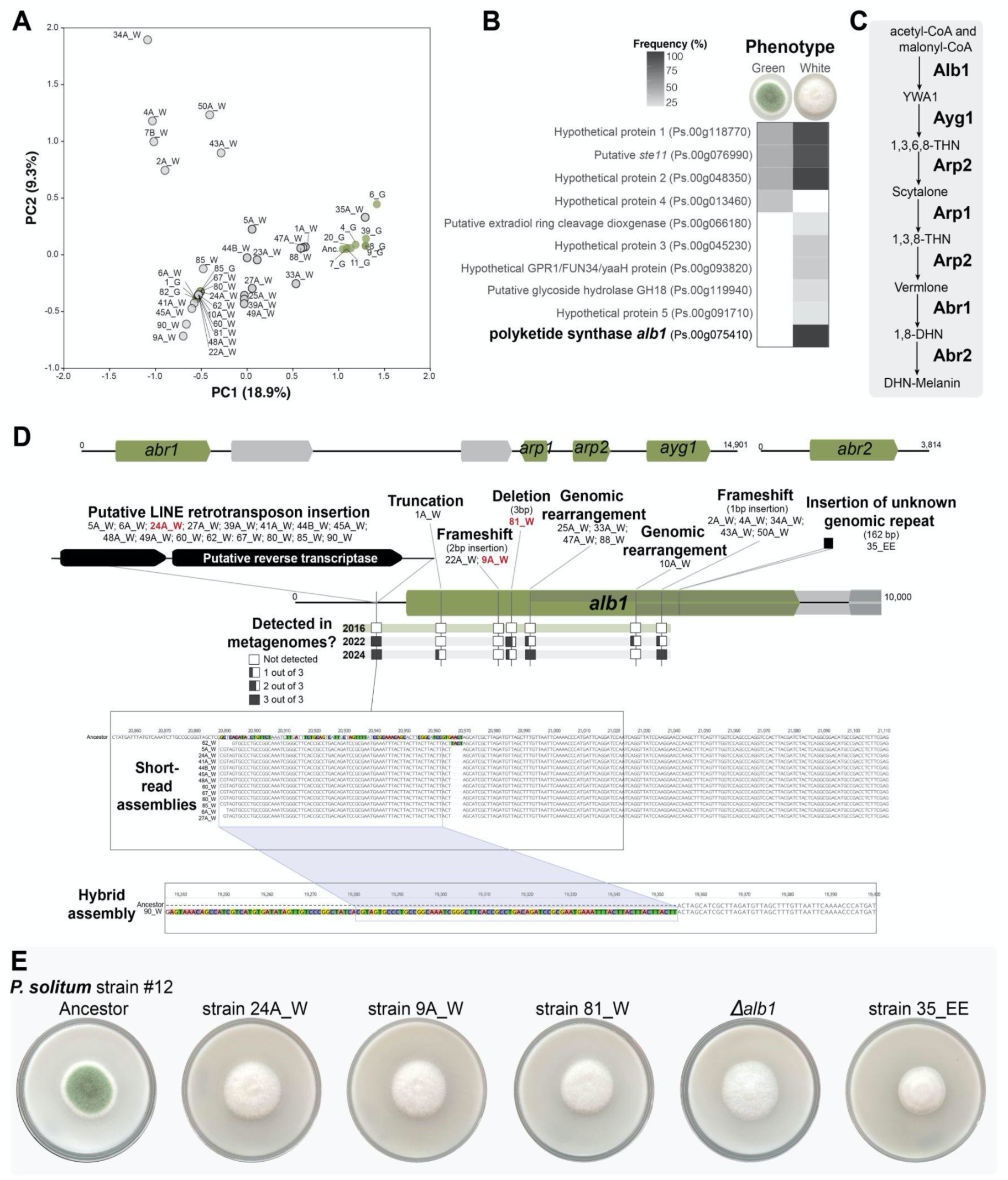
White *Penicillium solitum* strains have diverse mutations in the *alb1* gene. **(A)** Principal component analysis (PCA) plot of all detected SNPs (both synonymous and non-synonymous) across both green and white *P. solitum* strains. **(B)** Heatmap showing the frequency of non-synonymous mutations detected more than once across all *P. solitum* strains. Frequencies are calculated as a proportion of all isolates within each phenotype (green or white). Labels in parentheses are gene IDs in our reference genome assembly. **(C)** DHN-melanin biosynthesis pathway in *Penicillium* species (adapted from ^36^). **(D)** Location and type of mutations in the *alb1* gene, with other genes in the melanin biosynthesis pathway shown for context. Numbers under descriptions of types of mutations indicate the names of strains with those mutations. Strains highlighted in red were used for additional experiments below. Boxes with various amounts of fill indicate how often different mutations were detected in metagenomes sampled in 2016, 2022, and 2024 (see Figure 1E). **(E)** Phenotypes of various *P. solitum* strains, including the ancestor (left) and white phenotypes (strains 24A_W, 9A_W, and 81_W), knockout mutant (***Δ**alb1)*, and an experimentally evolved white phenotype (35_EE).

To identify mutations that could explain the phenotypic shifts in this *P. solitum* population, we searched for non-synonymous mutations present in all white isolates but absent in all green isolates (**Figure 2A; Table S4**). Only one genomic region matched these criteria: the polyketide synthase gene *alb1* that is part of the DHN-melanin biosynthesis pathway (**Figure 2B-C**). All white strains had either point mutations or large deletions within *alb1* or large insertions in the promoter region, with each strain only having one type of mutation in *alb1* (**Figure 2D; Figure S4-7**). Mapping of metagenomic reads from 2016, 2022, and 2024 (from Figure 1D above) confirmed that mutations were not present or below our detection levels in 2016, but a range of *alb1* mutations were observed in 2022 and 2024 with the promoter insertion mutations being the most common (**Figure 2D**).

Fungal melanins derived from polyketide pathways (e.g. DHN-melanin) are either black, brown, gray, or green pigments that protect fungi from UV radiation and other stressors ^34^. In *Penicillium* spp., melanins are predominantly constructed of dihydroxynaphthalene (DHN) ^35,36^. Based on studies in *Aspergillus* and *Penicillium*, the DHN-melanin pathway starts with the polyketide synthase (PKS) encoded by *alb1* (**Figure 2C**). This PKS converts acetyl-CoA and malonyl-CoA into the naphthopyrone compound YWA1, which is then followed by several enzymatic reactions that lead to the formation of melanin ^36,37^ (**Figure 2C**). The genes encoding the components of the melanin biosynthesis pathway are often found in biosynthetic gene clusters in ascomycete fungi ^38^, and are also clustered across three regions in our *P. solitum* genome (**Figure 2D**).

The most common alteration to the *alb1* genomic region was a large insertion upstream of the gene, detected in half of the sequenced white *P. solitum* strains (**Figure S4**). A hybrid assembly (Illumina short-reads and Oxford Nanopore long-reads) of one white strain (90_W) allowed us to identify this region as a putative transposable element (TE) in the long interspersed nuclear element (LINE) class of retrotransposons based on 100% DNA sequence similarity with a LINE TE annotated by RepeatModeler (**Figure 2D**, **Figure S4**). This 5,795 bp TE had a copy number of 25 and 19 in the white strain 90_W and the ancestor, respectively, suggesting a TE expansion in 90_W. Analysis of the TE genomic locations indicated that 10 copies in the white *P. solitum* strain 90_W were in different locations relative to the Ancestor assembly, including in the *alb1* upstream region (**Table S5**). In fungi, LINEs are typically 1-6 kbp in length and can disrupt regulation of neighboring genes within a 1 kbp range ^39^. Retrotransposons need an RNA intermediary in its transposition cycle ^40^ and the gene directly upstream of *alb1* (3.9 kbp length) in this TE cluster is highly similar to putative reverse transcriptases in other fungal genomes (95% percent DNA sequence similarity and 52% query coverage BLAST match with a predicted reverse transcriptase from *P. chrysogenum*, NCBI accession XM_066710680.1; **Figure 1D**). TEs can lead to adaptive genetic variation within species and populations ^41–43^ and when inserted in a regulator of a polyketide synthase gene similar to *alb1*, it can result in a decrease in melanization in a fungal wheat pathogen ^44^. Given this TE’s position and variation in copy number, it is likely a key driver in regulating expression of the *alb1* gene and consequently, the white phenotype.

To confirm whether disruption of the *alb1* gene leads to a white phenotype in *P. solitum* as observed in other *Penicillium* species ^36^, we knocked out the *alb1* gene by constructing a gene deletion cassette that replaced the *alb1* gene with a hygromycin resistance marker (**Figure S8**). Disrupting *alb1* resulted in a complete loss of green pigmentation and a colony morphology that closely resembles the white cave isolates (**Figure 2E**). We do not know if all steps of the DHN-melanin pathway previously identified in other *Penicillium* or *Aspergillus* species are identical in *P. solitum*, but this knockout strain demonstrates that the *alb1* gene plays a critical role in the distinct green pigmentation of this fungal species.

To experimentally demonstrate that white *alb1* phenotypes can appear in initially green populations of *P. solitum*, we performed experimental evolution of *P. solitum* strains through a serial passaging protocol as outlined previously ^28^. We serially transferred eight replicate populations of five strains (n=40) onto cheese curd agar (CCA) weekly, for 26 weeks. White colonies emerged at week six of the experiment, but stayed at a very low frequency (<5%) throughout the duration of the experiment (**Figure S9**). At the termination of the experiment (week 26), we sequenced the genome of a white isolate that looked very similar to those we observed in the cave and observed a 162 bp insertion in the *alb1* gene (**Figure 2D**). This sequence is repeated six times throughout the genome in intergenic regions, suggesting it could be a transposable element or other type of mobile DNA, but it does not match to sequences in databases of known transposable elements. These experimental evolution results demonstrate that loss of pigmentation via disruption of *alb1* is possible over short time scales in initially green *P. solitum* populations.

### White phenotypes have reduced melanin and spore production and altered global gene expression

Past comparative studies of the cheese mold *P. camemberti* found that domesticated strains had shifts in multiple traits beyond pigment production, including reductions in spore production ^7^. To characterize phenotypic traits that changed in the white *P. solitum* strains, we first confirmed that the observed pigment shift from green to white is associated with shifts in melanin quantities. There was significantly lower melanin production in all white strains compared to the green ancestor strain (F_4,35_ = 7.82, *p* < 0.001; **Figure 3A**). We also quantified the abundance of spores across several strains and observed similar reductions in spore production (F_4,15_ = 9.559, *p* < 0.0001), with all white strains producing significantly fewer spores (strains 24A_W, *p* = 0.0127; 81_W, *p* <0.001; 9A = <0.001; ***Δ**alb1*, *p* = 0.034) compared to the ancestor (**Figure 3B**). Unlike shifts in melanin and spore production, the only significant difference in mycelial growth was between two strains 24A_W and 81W (F_4,15_ = 6.155, *p* =0.004) and we did not see major changes in mycelial growth between the ancestor and white strains (*p* =0.004; strains 24A_W, *p* =0.281 ; 81_W, *p* =0.152; 9A =0.996; ***Δ**alb1*, *p* = 0.429; **Figure 3C**). These phenotypic shifts that occurred in a period of several years in this *P. solitum* population align with the reduced investment in costly biological traits observed in past comparative studies of cheese fungi ^7^.

**Figure 3:**
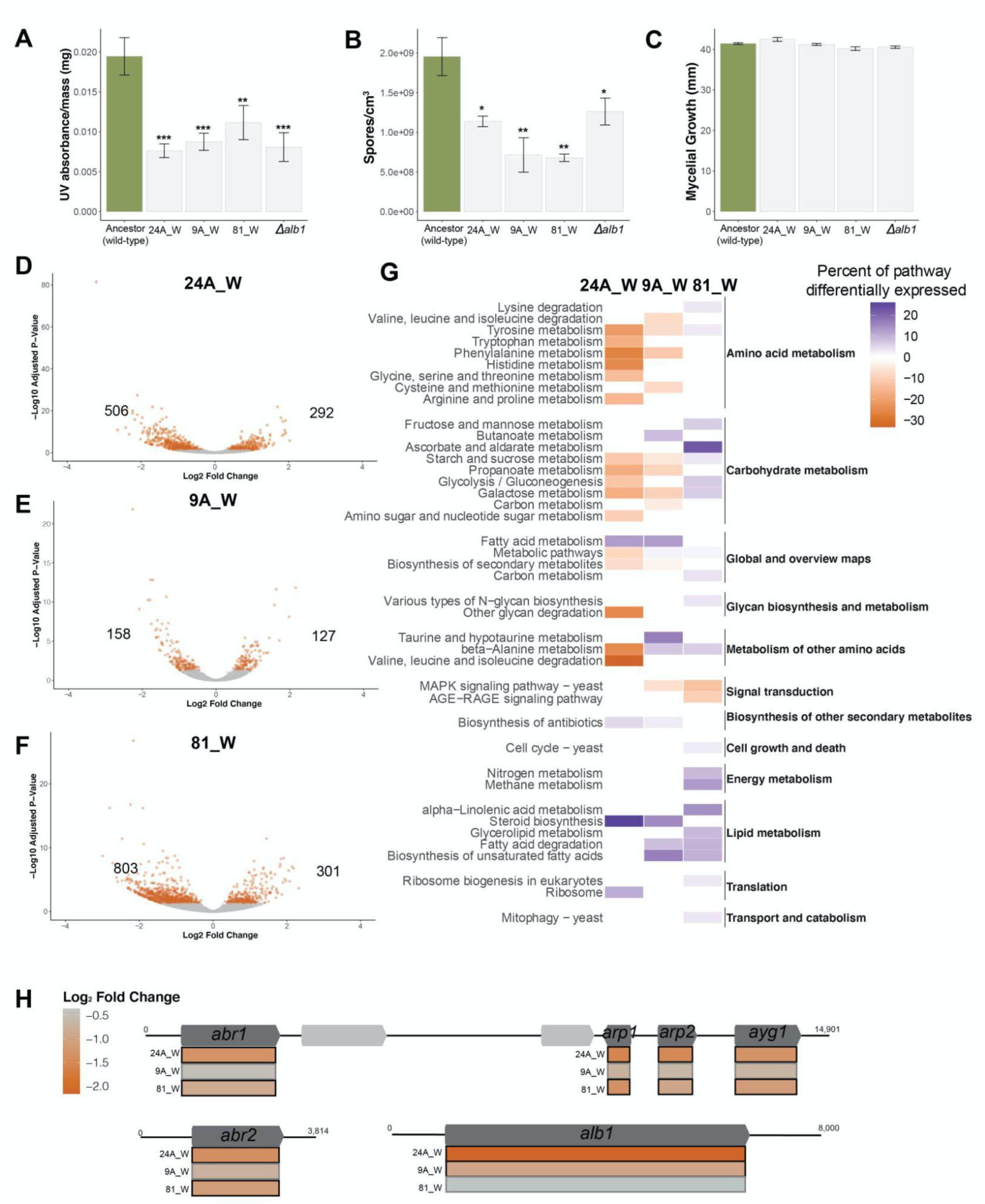
Phenotypic and transcriptomic shifts in white strains of *Penicillium solitum*. **(A)** Spore production per area of mycelium of ancestor and white phenotypes. Bars represent mean spores per unit area across four replicates. Error bars are one standard deviation of the mean. **(B)** Relative melanin production of ancestor and white phenotypes. Error bars represent means and one standard deviation calculated from eight technical replicates recorded from two separate experimental replicates (four technical replicates per experiment). **(C)** Mean mycelial growth of ancestor and white phenotypes. Error bars are one standard deviation of the mean. In A-C, asterisks indicate differences between white strains and the ancestor (*p*-values from Dunnett’s test multiple comparison) where * = *p* < 0.05, ** = *p* < 0.01, and *** = *p* < 0.001. **(D-F)** Volcano plots illustrate overall differences in gene expression between white phenotypes and the ancestral strain, with each dot representing an individual gene. Orange dots represent genes that are significantly differentially expressed (P < 0.05). Numbers indicate the total number of genes that are decreased in expression (left) and increased in expression (right) in each strain relative to the ancestor. **(G)** Heatmap displaying KEGG pathways (y-axis) significantly up-(blue) or downregulated (orange), by strain (x-axis). More detailed pathways (left) are grouped by broader gene pathway categories (right). **(H)** Fold change in gene expression of six genes in the DHN-melanin pathway. Genes are ordered according to position in the genome, and scaled according to size. Specific genes that are significantly downregulated have bolded borders.

To understand other global changes in the biology of cave-adapted white strains compared to the ancestral green strains, we measured global changes in mRNA expression by performing RNA sequencing (RNA-seq). We used the same green ancestral strain used as a reference above, as well as three white strains with unique *alb1* mutations (strains 24A_W, 9A_W, and 81_W) for RNA-seq. All three white strains showed shifts in global gene expression compared to the ancestor (**Figure 3D-F; Table S6-S8**). The proportion of significantly differentially expressed genes (DEGs) relative to the total number of genes (12,193) in the *P. solitum* genome, ranged from seven, two, and nine percent for strain 24A_W, 9A_W, and 81_W, respectively. Across all three strains, more DEGs were downregulated than upregulated, which is consistent with some previous reports of downregulation in global gene expression during adaptation via loss of function ^28^.

While a common pattern across all white phenotypes of *P. solitum* was an overall downregulation of genes, we did not see a consistent set of biological functions represented in the differentially expressed genes. Each strain had a unique set of pathways that were up- or down-regulated. For example, in strain 24A_W, five pathways relating to amino acid metabolism and one relating to carbohydrate metabolism were downregulated. These included valine, leucine and isoleucine degradation, phenylalanine metabolism, tyrosine metabolism, tryptophan metabolism, and galactose metabolism. There was a shared signal in the upregulation of genes in ergosterol biosynthesis (**Figure 3G**). Collectively, these data suggest that different white phenotypes of *P. solitum* that diversified in this cave environment have diverged in their underlying biology.

To directly relate expression data to mutations potentially underlying phenotypic changes, we examined differential expression of the six individual genes in the DHN-melanin pathway in *Penicillium* ^36^. In Strain 24A_W, all the genes in the pathway were significantly downregulated (**Figure 3H**), which could likely be explained by the insertion of a putative TE in the promoter region of *alb1* (**Figure 2D)** in this strain. In strain 9A_W, the *alb1* gene was significantly downregulated, but not the subsequent genes in the pathway. Conversely, in strain 81_W, all the genes in the pathway were downregulated, except for the *alb1* gene itself.

### White *Penicillium solitum* strains can outcompete green strains, but only in the dark

Our data above suggested that white, cave-adapted strains of *P. solitum* have outcompeted the ancestral green *P. solitum*, have accumulated mutations in a key gene in the melanin biosynthesis pathway, and have reduced expression of genes in the melanin biosynthesis pathway. Melanin is thought to be important for protecting fungi from stress in natural environments with high light levels ^45,46^ and may be an unnecessary cost in environments with lower light levels such as cheese caves. Loss of melanin production is a hallmark of adaptation to cave environments in multiple domains of life ^47,48^, and may also be a key evolutionary shift during adaptation of fungi to dark cheese aging environments. We predicted that the success of the *P. solitum* white strains in this population was due to a higher fitness advantage from reduced production of costly melanin and downregulation of other biological processes observed in the RNA-seq data.

To test this prediction, we performed pairwise competition experiments on cheese curd to determine whether white phenotypes would outcompete green phenotypes and whether this was dependent on light (**Figure 4A**). We competed several white strains (24A_W, 81_W, and Δ*alb1*) in one-to-one ratios with the ancestral green strain and measured abundances of each strain after seven days of growth. Experiments were conducted in both the dark and in the presence of full spectrum light that mimicked light intensity on a sunny day in New England (see Methods). In the dark, the two white strains from the cave (24A_W, 81_W) outcompeted green strains (**Figure 4B**), aligning with their ability to displace the green strain in the cave over time. Interestingly, the Δ*alb1* strain was not able to outcompete the ancestor in the dark, suggesting that the complete loss of *alb*1 activity was not sufficient to give this strain a competitive advantage, or other mutations in the genomes of 24A_W and 81_W may confer a competitive ability over the ancestor. Light had an overall significant effect on competitive ability (F_1,39_ = 26.76, *p* < 0.00001), but that effect was significantly dependent on strain type (F_2,39_ = 4.34, *p* < 0.05). The presence of light reduced or completely flipped the competitive advantage of 24A_W and 81_W. Surprisingly, there was no significant difference in competitive ability between light and dark treatments for the *alb1* knockout strain (**Figure 4B**). These results confirm our prediction that cave-adapted strains have a fitness advantage over green strains, and that this advantage is dependent on light conditions.

**Figure 4:**
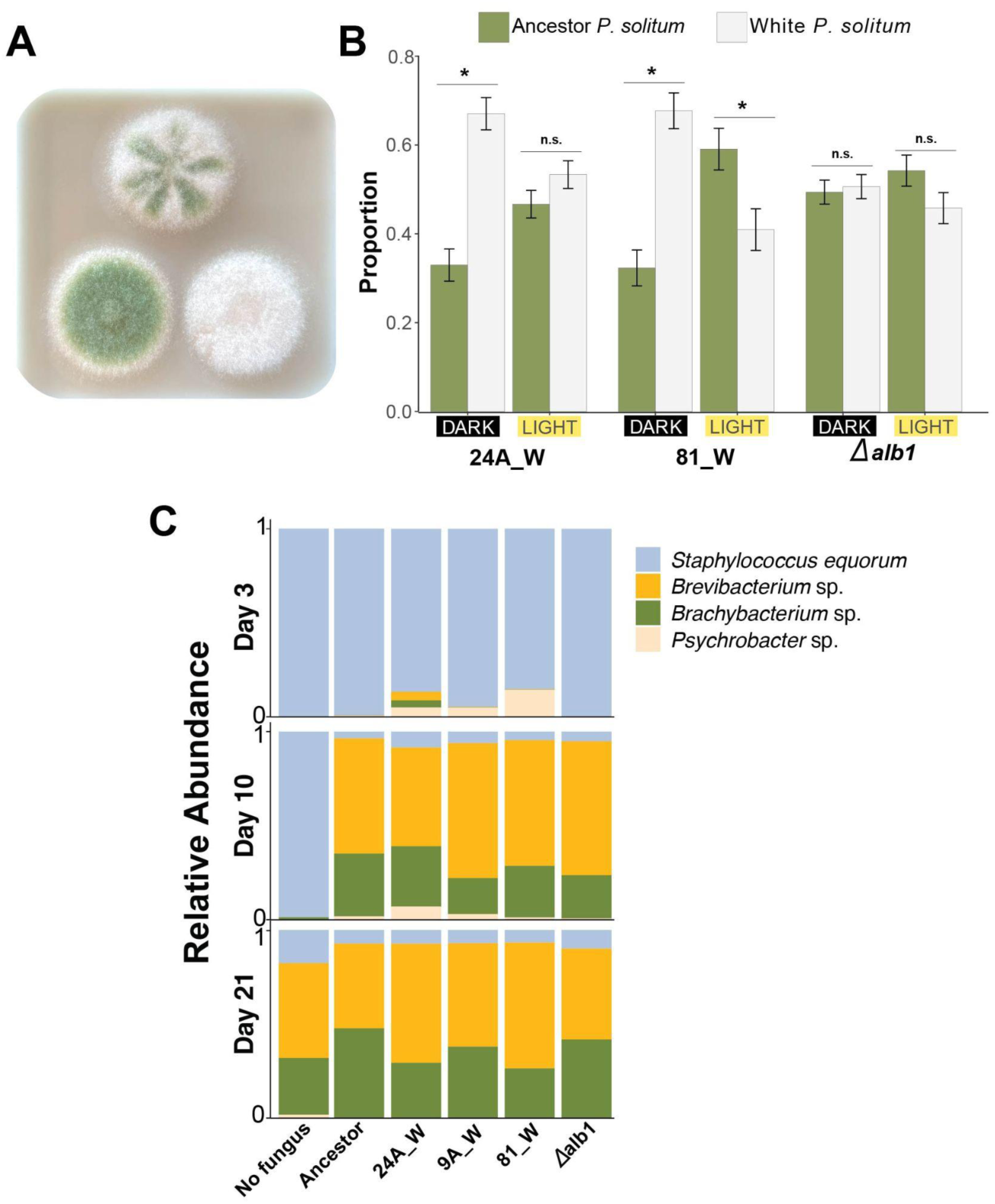
Consequences of cheese cave adaptation in *P. solitum*. (**A)** Photographs of a mixed culture (center, top) and monocultures (ancestor, bottom and left; 24A_W bottom and right) from the dark treatment. **(B)** Proportion of CFUs of ancestor (green) and various white strains (white) in co-culture in light and dark treatments. Bars are calculated as the mean proportion of green and white (both bars per treatment add up to one) colonies (CFUs/mL) across eight technical replicates from two separate experiments (four replicates per experiment). Error bars denote one standard deviation from the mean. * = significant (p < 0.05) difference in CFUs between green and white strains based on t-test. n.s = not significant. **(C)** Relative abundance of four bacterial species when growing in a community alone (“No fung.”) or in co-culture with various white strains at days three (top), 10 (middle) and 21 (bottom). “Anc.” = ancestor.

### Phenotypic shifts in *Penicillium solitum* strains do not have major consequences for bacterial community assembly

On cheese rinds, *Penicillium* species mediate bacterial community assembly dynamics through the production of antibiotics ^49^, volatiles ^50^, by modifying resource availability ^32^, or changing the physical substrate to aid in dispersal ^51^. Since novel phenotypes show altered global gene expression and decreased melanin production compared to ancestor strains, we predicted that short-term phenotypic shifts would have consequences for how cheese rind bacterial communities assemble. To test this, we assembled a four-member bacterial community that includes some of the most widespread taxa of cheese rinds ^25^. We quantified community assembly dynamics through changes in bacterial alpha diversity and community composition by measuring total CFUs of each community member at days three, 10, and 21, grown without a fungus, with the ancestor *P. solitum,* the *alb1* knockout, and the same three white strains used in RNA-seq. Many cheese rind bacteria depend on free amino acids produced through the proteolytic activities of *Penicillium* and other fungi in cheese rind communities ^32,52,53^. To quantify consequences of changes in ecologically relevant enzymatic activities, we compared the release of free amino acids from casein degradation between ancestor and cave-adapted *P. solitum* strains. Past studies have suggested that domestication of molds in food production environments can increase their extracellular enzyme activities, leading to increased decomposition of food substrates ^54–57^. We predicted that white strains of *P. solitum* may have increased free amino acid concentrations as they adapt to the cheese environment.

Surprisingly, there was very little difference in bacterial community assembly between green and white strains (**Figure 4C, Figure S10-S11**). The only case where alpha diversity differed between strains was on day three, where it was higher in strain 24A_W than in the Ancestor (**Figure S6**). Similar to patterns in alpha diversity, a permutational analysis of variance (PERMANOVA) found only marginally significant differences in community composition between communities assembled with strain 24A_W and the ancestor on day three (**Figure S7**; *p* = 0.030), with no significant differences in community composition between mutants and the ancestor on days three and 21 (P > 0.05). We also saw very little effect of strain identity on the release of free amino acids from cheese curd (**Figure S12;** *p* = 0.245), and this may explain the muted impacts of strain identity on bacterial community composition. These data suggest that while the green to white phenotypic shift can have major impacts on the biology of *P. solitum*, these changes do not translate to shifts in how cheese rind communities form in this environment.

## DISCUSSION

Melanin allows fungi to survive in variable environments by increasing cell wall rigidity and their ability to cope with environmental stressors like UV radiation, temperature, and high levels of salinity ^44,58–61^. A previous study on an endophytic fungal species found positive selection on a genomic island that contains the DHN-melanin biosynthetic gene cluster at higher salinity levels ^62^. Others have found that variation in melanin accumulation among strains of a fungal wheat pathogen could be explained by insertions of transposable elements in the DHN-melanin biosynthetic gene cluster ^44^. These past comparative studies have highlighted the variable nature of melanin production within fungal species, but do not illustrate exactly how quickly and how melanin biosynthesis genes evolve within populations.

Our study provides evidence of *in situ* adaptation of a fungal population, and suggests that melanin and other traits can rapidly evolve in dark food production environments. An increase in white phenotypes of *P. solitum* compared to green corresponded with mutations found exclusively in white strains on or in the promoter region of the *alb1* gene. By recreating the cheese cave environment *in vitro*, we were able to confirm our hypothesis that melanin production comes at a fitness cost in the absence of environmental stressors like light. These results suggest that white strains may be adapted to dark cheese cave conditions. This loss of pigmentation during adaptation to an artificial cave is similar to how a range of organisms adapt to natural cave environments ^63–65^.

This work also adds to a growing body of papers showing how TEs may facilitate fungal adaptation ^39,66–70^ by causing changes in expression of neighboring genes. Our findings align with a recent study that attributed variation in melanin accumulation of a fungal wheat pathogen to an insertion of transposable elements upstream of a promoter that regulates gene expression in the polyketide synthase 1 gene cluster ^44^. The expansions of TEs in genomes have been shown to promote the domestication of crop plants ^41,71,72^, but little is known about how TEs enable the domestication of fungi. A recent study showed that domesticated fungi contain a higher TE content compared to closely-related wild relatives ^73^, but this work did not determine how TE expansions facilitated past domestication events. To our knowledge, our work is the first case demonstrating how TEs can directly facilitate fungal domestication in a food environment. TE insertions can cause genome instability and a TE insertion may not be permanent^74^. Future studies with this system should determine whether TE insertions that disrupt *alb1* expression are stable and maintain the green to white phenotypic shift.

Comparative genomic studies have demonstrated that several *Penicillium* species have been domesticated by humans ^75^. *Penicillium roqueforti,* which is used to produce blue cheese, was likely domesticated multiple times ^24^. Similarly, the ancestor of *P. camemberti* used to age Brie and Camembert, underwent two consecutive domestication events transitioning from a blue-green phenotype to a white fluffy phenotype that morphologically resembles the cave-adapted strains in this study ^7^. In both mentioned cases, the hallmarks of fungal domestication to a cheese cave environment are broadly defined by an increase in expression of traits beneficial in cheese production (increased lipolysis, proteolysis, etc.) at the expense of traits adaptive to variable natural environments (melanin, spore, and/or mycotoxin production) ^76^. Our data from this one *P. solitum* population are consistent with these previous studies showing a decrease in pigmentation and spore production.

One unique aspect of our temporal sampling of the same cheese and fungal population is it provides a window into the process of unintentional microbial domestication. We know that many food fungi have been domesticated, but we do not know *how* humans selected strains for use in food and how they initially found these strains ^77^. Our work demonstrates that strains with desirable properties for cheese production can take over populations of cheese-associated fungi without any direct or intentional inoculation or manipulation from the food producers. The green-gray rinds that were observed during the early stages of the *P. solitum* colonization of this blue cheese were perfectly acceptable. But the shift to a rind dominated by white strains of *P. solitum* is considered a positive outcome by the cheese producers (one of the co-authors of this work is a producer of this cheese), because cheesemakers often need to manage rind fungi to suppress green and blue molds for consumer acceptance purposes ^26^. Understanding how cheese strains diversify within a single production facility may allow unintentional domestication to become more deliberate. For example, manipulating light conditions in caves may help select for specific strains of *Penicillium* based on their fitness in different light environments.

Our work provides insight into the timing and molecular mechanisms of fungal adaptation in cheese production environments. Beyond aesthetics, the impacts of *P. solitum* evolution on the flavor properties of this cheese are not yet known. Our free amino acid data suggest that flavor development due to proteolysis may not be impacted, but other processes such as lipolysis, production of volatile compounds, and protection of cheese from invasion by pathogens and spoilage organisms should be considered in future studies. Most work on the domestication of filamentous fungi in food production has focused on inferred past domestication events in European or Asian foods ^7,55,75,76,78,79^. Most American artisan foods, including surface-ripened cheeses, are made with descendants of these European and Asian molds. Our work in an American cheese cave suggests that domestication of food fungi can occur outside of traditional production regions and may provide a source of local cheese cultures for American fermented food production. We predict that transposon-mediated rapid trait evolution can occur in many food-associated fungal species in production and aging facilities around the world. Future work to characterize the causes and consequences of fungal adaptation across different cheese cave environments can help identify strains with potential as novel starter cultures for cheese production.

## RESOURCE AVAILABILITY

### Lead contact

Requests for additional information and resources should be directed to the lead contact, Benjamin Wolfe.

### Materials availability

Strains of the fungi used in this study are available from the lead contact upon request.

### Data and code availability

Raw shotgun metagenomic reads have been submitted to NCBI with accession numbers indicated in the supplementary information. Raw sequence reads from various *P. solitum* strains have been submitted to the NCBI under accession numbers provided in the supplementary information. Hybrid genome assemblies of the *P. solitum* ancestor strain and white strain 90_W have been submitted to NCBI with Biosample accession numbers SAMN47278713 and SAMN47278714, respectively. All other data reported and code made to analyze data in this study will be made available upon request to the lead contact.

## ACKNOWLEDGEMENTS

We are indebted to the many cheese producers and staff at Jasper Hill Farm and the Cellars at Jasper Hill that have worked to make and age Bayley Hazen Blue. This work was supported by the National Science Foundation (CAREER IOS/BIO 1942063) to B.E.W and the National Institutes of Health (3R01GM112739-07W1) to J.L.E. We are very grateful to Charlie Kalish for proposing to Rachel Dutton in 2016 at Jasper Hill. It is because of his marriage proposal that the 2016 samples were collected and provided a valuable time capsule for this project.

## AUTHOR CONTRIBUTIONS

N.L.L. and B.E.W conceived the project. N.L.L., B.E.W, and J.L.E. performed lab work. N.L.L., B.E.W, and J.L. conducted bioinformatic analyses. N.L.L. and B.E.W prepared figures and tables. N.L.L., B.E.W, and M.K. coordinated sampling and access to the cheese facility. N.L.L. and B.E.W drafted the manuscript and all authors contributed to the editing.

## SUPPLEMENTAL INFORMATION

See Tables S1-S10 and Figures S1-S12.

## MATERIALS AND METHODS

### Study site description and cheese rind sampling

Cheese rind samples were collected from the Cellars at Jasper Hill in Greensboro, VT, USA. Sampling took place in the summers of 2016, 2022, and 2024, within the same vault, where the blue cheese Bayley Hazen Blue is aged. Within this vault, 50 different wheels were sampled across a range of cheeses of different ages, from within a week of being produced to over two months of aging.

To collect samples, we followed methods outlined previously ^25^. Briefly, we scraped the surface of the rind of each cheese with sterile wooden dowels to obtain approximately 100-200 mg of material. We stored the scraped contents in both 15% glycerol in 1X phosphate buffered saline (PBS) (for plating out and isolating fungi) and molecular grade water (for metagenomic sequencing). We stored rind samples at −80°C until further processing.

We isolated abundant fungi from these collections through serial dilutions of the glycerol samples onto plate count agar supplemented with 0.1% whole milk powder and 1% salt (PCAMS^80^) with chloramphenicol (50 μg/ml). To count the proportion of white versus green *P. solitum* colonies, we morphologically identified colonies as *P. solitum* and counted the total number of white and green colonies from each output plate representing a cheese rind sample. For the purpose of comparing proportional phenotypic changes over time, we excluded samples with output plates that contained less than five total CFUs of *P. solitum*, which included four samples from 2024 and four from 2022. For the remaining 142 samples, we calculated the proportion of white versus green *P. solitum* as the fraction of white to total number of *P. solitum* CFUs within each year (**Figure 1E)**.

We selected a proportion of phenotypes that were representative of that year for whole-genome sequencing. For example, in 2024 we only selected white colonies because it was very difficult to find green colonies, in 2022 we selected eight white colonies and two green ones, and in 2016, we selected eight green colonies. We sequenced more isolates from 2024 than previous years to confirm the robustness of genetic patterns we observed between 2016 and 2022.

### Shotgun metagenomic sequencing

For shotgun metagenomic sequencing, we selected three rind samples from three separate wheels of cheese from each year. Samples were chosen from the oldest (∼60 days of aging, in the later stages of rind succession for this cheese^25^) cheese wheels per year, to standardize for variation in cheese rind community succession ^25^. We extracted DNA using the Qiagen Dneasy Powersoil Pro Kit following manufacturer’s details ^81^ by taking a 250mg subset of homogenized cheese rind samples. Shotgun metagenomic libraries were prepared and sequenced on the Illumina NextSeq2000 platform with 2 x 150 bp at SeqCoast Genomics in Portsmouth, NH. Read demultiplexing, trimming, and analytics were carried out with DRAGEN v3.10.12.

To identify the most abundant taxa, we classified the paired-end raw reads (ranging from ∼500,000 - ∼2,000,000 reads per sample) using Kaiju (v1.9.0) ^82^. Kaiju parameters were set to greedy mode that assigns reads based on its alignment against the NCBI reference database, including eukaryotes and non-redundant sequences. The allowed mismatches and match scores were kept at default values. To visualize communities over time, the classification results were filtered to species-level resolution, displaying only taxa that exceeded the default abundance threshold set in Kbase ^83^ visualizations. For visualization, we separately plotted metagenomic communities with and without reads assignment as “unclassified”. The proportion of unclassified reads averaged by year were 61, 55, and 52% for years 2016, 2022 and 2024, respectively (**Figure S2).**

### Whole-genome sequencing and variant identification

To select colonies for whole-genome sequencing, we made pure isolation streaks of single colonies from 10 samples in both 2016 and 2022, and 29 in 2024. Across all years, we selected colonies that were isolated from unique cheese wheels that were distributed throughout the aging room to fully capture rind samples at various stages of aging. Once purified streaks were mature after growing for seven days in the dark at 24°C on PCAMS, we harvested fungal tissue which we stored in molecular grade water at −80°C until further processing.

DNA for whole-genome sequencing was prepared using the same methods described above. At SeqCoast Genomics (Portsmouth, NH, USA), each strain was sequenced with 15-20X coverage with paired-end 150bp libraries on an Illumina NextSeq2000 platform. Read demultiplexing, trimming, and run analytics were performed with DRAGEN v4.2.7, a built-in software on the NextSeq2000 platform.

To create a complete draft assembly of the green reference *P. solitum* strain (referred to as “#12” or “ancestor” throughout the manuscript), we performed both short (using methods above) and long-read sequencing on an Oxford Nanopore platform at SeqCoast. For long-read sequencing, DNA was extracted with the DNAeasy 96 Powersoil Pro Qiacube HT Kit. Cell lysis was performed with MaxMAX Microbiome bead beating tubes, with an additional lysis step included by incubating cells in 20U of Zymo’s Zymolase Ultra for one hour at 37°C. Samples were prepared with Oxford Nanopore Technologies SQK-NBD114 native barcoding kit. Long fragment buffer was used to promote longer read lengths, with sequencing performed on the GridION platform, using a FLOW-MIN114 Spot-ON Flow Cell, R10 version with a translocation speed of 400 bps.

We assembled the Nanopore reads with Canu v2.2 ^84^ using default parameters. This assembly was polished for two rounds with Pilon v1.24 ^85^. Raw illumina reads were aligned to the Canu-assembled genome using BWA-MEM ^86^ and resulting alignments were processed using SAMtools ^87^ to generate inputs for Pilon v1.24 ^85^ to perform error correction, with default settings. We annotated the hybrid assembly in the GenSAS ^88^ workspace using Augustus ^89^ as a gene model prediction tool.

To detect variants of cave isolates, we mapped raw reads to the reference genome with Geneious’s read mapper (Version 2021.2.2) set to medium-low sensitivity, minimum quality to 30, maximum gap size to 15, minimum overlap identity 80%, and maximum ambiguity to four. Variants were called with Geneious Variant Caller using conditions of a minimum coverage of 20X and variant frequency of 90%. To determine the putative functions of genes with non-synonymous mutations (**Figure 2B**) that were not clearly annotated in our genomes, we used BLASTp searches against the NCBI non-redundant protein sequences database. To visualize similarities in all SNPs among different strains, we created a SNP matrix of 92 SNPs detected across all green and white strains. Presence of rare alleles was coded as one and abundant alleles as zero ^90^ and then this matrix was used for a principal coordinate analysis (Past 4.01).

To determine potential structural variants in the *alb1* genomic region, we used alignments of whole-genomes assembled from paired-end Illumina reads. Genomes were assembled using the SPAdes genome assembler (v 3.15.3). We next aligned *alb1* sequences (including approximately 15 kbp upstream and downstream of *alb1*) from assembled genomes against the hybrid reference genome assembly using MUSCLE v3.8.425 ^91^.

To determine whether DNA insertions in the *alb1* region were putative transposable elements, we searched for exact sequence matches from outputs of RepeatModeler ^92^ generated during genome annotation in GenSAS.

To identify the presence of various *alb1* mutant sequences in the 2016, 2022, and 2024 metagenomes, we mapped shotgun metagenomic reads using the same approach described above for variant identification, but with the following settings: minimum overlap identity 98% (to avoid mapping reads of other *Penicillium* species), 10% maximum gap size per read (to allow for deletions), and maximum ambiguity of four. To determine the presence of SNPs and short indels, we mapped reads to the full length *alb1* gene of the reference genome. To detect presence of structural variants, we mapped reads to junctions that formed from TE insertions or genomic rearrangements. Because we were using 150 bp length reads, we used a mapping region that spanned 75 bp downstream and upstream of the junction site. Mutant sequences were considered present in a sample if metagenomic reads spanned the junction 10 bp upstream and downstream of the junction.

### RNA extraction, sequencing, and analysis

To generate fungal cultures for RNA-seq, we inoculated three different *P. solitum* strains onto CCA. For the strains, we used the same ancestor (#12) as we did for read mapping and variant calling, along with three cave-evolved white strains (81_W, 9A_W and 24A_W). We selected three different white strains for RNA-seq to represent the breadth of mutations that we observed in *P. solitum*. Strain 81_W was isolated from 2022 and represented the only strain from that year with a point mutation. We selected strain 24A_W, because it has a putative TE insertion in the promoter region of *alb1*, which was the most common type of *alb1* variant in our dataset. Strain 9A14 had an insertion (+GT) that caused a frameshift.

We inoculated three spots (200 CFUs per spot) evenly spaced across the 20 ml of CCA in a 100-by-15 mm Petri dish per each of the four experimental replicates (unique plates per strain). After five days of incubation at 24°C in the dark, we excised fungal tissue from CCA using sterile surgical blades. We stored the harvested fungal material in three ml of RNALater (Qiagen, Germany) at −80°C for downstream processing.

We thawed and extracted RNA from one spot per experimental replicate, per strain, using the Qiagen RNeasy Plant Mini Kit using previously described methods^28^. We performed this extraction after grinding the sample submerged in liquid nitrogen with sterile mortars and pestles. Using this approach, we placed approximately 100 mg of the ground fungal tissue in 750 µl of RLT buffer (Qiagen) with 10 µl 2-mercaptoethanol per one ml added to the RLT buffer. For the subsequent extraction steps, we followed the protocol as outlined by the manufacturer’s recommendations. RNA preparation for differential expression comparison between strain 81_W and the reference strain was performed in a separate run prior to the comparison of 9A_W and 24A_W to the reference strain.

RNA-seq library preparation and sequencing was performed at SeqCoast Genomics using an Illumina Stranded mRNA Prep kit with IDT Illumina Unique Dual Indexes. Libraries were sequenced on the Illumina NextSeq2000 platform using a 300-cycle flow cell kit that resulted in paired-end sequencing with 150 reads per end (2×150bp). Read demultiplexing, trimming, and run analytics were conducted with DRAGEN v3.10.12.

Ancestor and white phenotype reads were mapped to the assembled and annotated reference genome used for variant calling (see above) with Geneious Prime Read Mapper (Version 2021.2.2) set to the medium-low sensitivity parameters. We calculated differential expression between the ancestor (#12) and white strains with DESeq2 ^93^. To be considered differentially expressed, expression had to be greater than double (log_2_ ratio > 1 or < 1 for an increase and decrease in expression, respectively) in the white, compared to the green phenotype, with a p-value < 0.05, adjusted according to the Benjamini-Hochberg procedure ^93^. To determine any biological pathways that were enriched in the differentially expressed gene lists, we used the KOBAS-i Gene-list Enrichment tool ^94^, using *Aspergillus fumigatus* as the background genome. DESeq and KOBAS outputs were exported and imported into Rstudio v2024.04.2+764 to visualise as scatterplots and heatmaps using ggplot ^95^.

### *alb1* gene knockouts

*Penicillium solitum alb1* was deleted using homologous recombination to replace the open reading frame with a recyclable hygromycin B-resistance conferring cassette ^96^. The homologous flanks were amplified from *P. solitum* genomic DNA. The hygromycin resistance cassette was amplified from plasmid pSK529 ^96^. Primers used for these products are listed in **Table S9**. PCR products were stitched together as previously described ^97^. The final construct was amplified with nested primers situated ∼500 bp internal to the homologous flanks’ ends (**Figure S8**), and purified with a G50 column (Cytiva, Marlborough, Massachusetts, USA) to prepare for transformation. *P. solitum* protoplasts were prepared as previously described ^97^.

For the transformation, 20 µl of the G50-purified *alb1* deletion construct (approximately 1 µg) was mixed with 80 µl STC buffer and 100 µl of protoplasts and left on ice for one hour. A negative control was the same condition with 20 µl of ddH_2_O instead. Then, 1.25 ml of PEG solution (180 mM PEG, 50 mM CaCl_2_, 50 mM Tris-HCl, pH 7.5) was added and gently mixed thoroughly to incubate at room temperature for 20 minutes. The PEG solution was then diluted by addition of five ml STC buffer, then 10 ml of liquid sorbitol minimal media (GMM supplemented with 1.2 M sorbitol) was added to recover the protoplasts overnight at 25°C, 250 rpm. Lastly, 35 ml of sorbitol minimal media top agar (1% agar) supplemented with 100 µg/ml hygromycin B was added to each overnight recovery suspension, inverted to mix, then plated onto sorbitol minimal media plates supplemented with 100 µg/ml hygromycin B.

White transformants were screened for *alb1* deletions using two primer pairs, with one primer for each pair in the hygromycin-resistance cassette and the other within the genome outside of the recombined region (**Table S9; Figure S8**). One transformant successfully provided both of these amplicons, confirming the white phenotype was connected to *alb1* deletion (**Figure S8**). For this one successful transformant, we used a combination of Illumina short-read and Oxford Nanopore long-read whole-genome sequencing as described above to confirm the correct deletion of *alb1* (**Figure S8**).

### Experimental Evolution

We used the same reference strain of *P. solitum #12* used above, in addition to three green strains isolated from 2016 (6_G, 4_G, 39_G) and one green strain isolated in 2022. We chose green strains based on unique nonsynonymous point mutations (**Table S3**) for experimental evolution. Each strain was grown in 150 µl CCA in 1.5 ml microcentrifuge tubes. We inoculated 200 CFUs (a volume of 10 µl at a concentration of 20 CFUs/µl of each strain) into eight separate replicate populations.

After incubating at 24°C for seven days in the dark, we added 500 µl of 15% glycerol in 1xPBS to each microcentrifuge tube, which we homogenized with sterile pestles. From this homogenate, we transferred 10 µl to a fresh microcentrifuge tube containing CCA. Each week, homogenates were stored at −80 °C as a fossil record, and every other week, we plated out a subset of the homogenate onto PCAMS at dilutions 10^−3^ and 10^−4^.

After seven days of incubation in the dark, we counted the total number of CFUs and recorded the proportion of colonies that resembled *alb1* white phenotypes. At week 26, we isolated a pure white colony experimentally evolved from the ancestral strain 39_G (**Table S2**). We sequenced the genome of this strain and identified mutations in it using methods outlined above for cave strains.

### Melanin abundance, spore production, and mycelial growth

To measure melanin, we used a quantitative colorimetric assay with an azure A dye solution ^98^ with the following modifications. The azure A dye solution was made by dissolving the dye in 0.1M HCl. The resulting dye was then diluted until a measurement 0.665 at 610 nm absorbance was recorded. After growing the same strains used for RNASeq and the *alb1* knockout strain for seven days on PCAMS, we harvested fungal tissue, and recorded the mass harvested. Fungal tissue was placed and homogenized with sterile micropestles into 500 µl of azure A dye, before adding an additional 500 of µl of dye. This solution, in addition to four blank replicates without any fungal tissue, was incubated at 24°C on a shaking incubator for 90 minutes, covered with aluminum foil. After incubation, we centrifuged the solution at 15000 rpm to separate the aqueous solution from fungal material. We then transferred 200 µl of the supernatant (avoiding any fungal material) to a 96-well plate.

We measured UV absorbance on a SpectraMax M3 microplate reader at a 610 nm wavelength. We took the difference in absorbance values between samples recorded in blank samples (samples incubated without fungal tissue) relative to samples incubated with fungal tissue, and divided it by the initial weight added to get a change in absorbance value by unit of mass. A standard curve is typically made for the azure A melanin assay^98,99^, but since we were only interested in the magnitude of change of white strains relative to its green ancestor strain, we did not generate a standard curve. We repeated this assay twice with four technical replicates per experimental round and reported the mean values calculated across replicates (eight total per treatment) from both experimental rounds. To compare differences in melanin among strains, we performed a one-way ANOVA in R, followed by Dunnett’s test for multiple comparisons in R ^100^.

For the same set of strains used to quantify melanin, we measured conidial spore production per unit area of mycelium, in addition to total mycelial growth. For both of these assays, we spot-plated a volume of 10 µl of a concentration of 20 CFUs/µl (200 CFUs total) of each strain with four technical replicates onto the center of a 100-by-15-mm petri dish containing 20 ml of CCA. After seven days of incubation in the dark at 24°C, we measured mycelial growth as the diameter of the colony using a digital caliper. Using the same set of samples, we used a cork borer with a 6 mm diameter to harvest a plug of fungal tissue from the center of the colony, which we suspended and homogenized in 200 µl of 1X PBS. The density of spores in this suspension was determined using a Petroff-Hausser Counting Chamber hemocytometer. For both mycelial growth and spore production, we used four technical replicates and one experimental replicate per assay and performed the same statistical tests used for comparing melanin absorption (see above).

### Competition experiments

For competition experiments, we inoculated 100 CFUs of green and white *P. solitum* strains onto 25 ml of CCA in square 100-by-100-mm petri dishes. In addition to mixed treatments, we inoculated 200 CFUs (using the same volume as above in sporulation assay) of green and white colonies in monoculture on separate spots on the same petri dishes (**Figure 4A**). Both light and dark treatments were kept in the same temperature-controlled incubator at 24°C, with dark-treated plates covered in aluminum foil. Using a factorial design, within each light treatment and strain used, we had four technical replicates, placed in the same bag. The experiment was repeated using the exact same experimental setup for two experimental replicates. For the light treatment, we installed three LED full spectrum grow lights. These lights have 104 red, 140 blue, and 6 white LEDs that result in a wavelength ratio lower, but in a similar range to an outside environment on a sunny day at 65.43 µmol/photons/m^2^/s. Petri dishes were stored lid-side up to face lights, covered, and sealed with clear plastic bags to avoid desiccation.

After seven days of incubation, we harvested the colonies with a 18 mm diameter cork borer from the center of each colony. This plug was placed in a sterile Whirl-PAK ® sampling bag (Fort Atkinson, WI, USA) and homogenized in a 5 ml volume of 15% glycerol in 1X PBS. We plated this homogenate out on PCAMS plates and after seven days of incubation at 24°C in the dark, we counted the number of green and white colonies per replicate and calculated the number of CFUs/ml. To compare differences in competitive ability between light treatments, we used the proportion of white colonies relative to green as a competitive index. We next performed a two-way ANOVA to test the main effects of light and strain, in addition to the interactive effects of light*strain on competitive ability. This was done by fitting a linear model (Competitive Index ∼ Light*Strain) and using Type two sum of squares (“Anova” in the ‘car’ package ^101^). We performed post-hoc pairwise comparisons with estimated marginal means ^100^ to test the differences in competitive ability by light treatments and light by strain interactions. To statistically compare the difference in CFUs between the two phenotypes by treatment and strain, we performed paired t-tests.

### *In vitro* community assembly

For the community assembly experiment, we inoculated 200 CFUs of each member of the bacterial community (*Staphylococcus equorum* BC9, *Brevibacterium auranticum* JB5, *Brachybacterium alimentarium* JB7, and *Psychrobacter sp.* strain JB193) either alone, or with *P. solitum #12* (ancestor), *Penicillium alb1* knockout*, P. solitum* strain 9A_W (cave isolate), *P. solitum* strain 24A_W (cave isolate), or *P. solitum* strain 81_W (cave isolate). For each treatment, we included four replicate 1.5 ml microcentrifuge tubes filled with 150 µl CCA for harvest days three, 10 and 21, to capture differences in community assembly throughout successional time. At each harvest time, we added 500 µl 15% glycerol in 1X PBS to the 1.5 ml microcentrifuge tubes, which we homogenized with sterile pestels. Homogenates were diluted and plated onto PCAMS with chloramphenicol (50 μg/ml) to count fungal growth in the absence of bacteria, and PCAMS supplemented with cycloheximide (100 mg/L) to count bacterial growth in the absence of fungi. Each bacterial community member has a different color and colony morphology which enabled us to quantify abundance by counting colonies and calculating CFUs/ml of each strain ^80^.

We calculated bacterial alpha diversity with the Shannon diversity index with the ‘Vegan’ package in R ^102^ and then tested the significance of differences in diversity between ancestor, cave-adapted strains, and the knockout strain with a factorial ANOVA, with the host treatment factored by harvest day. Next, we performed post-hoc multiple pairwise comparisons using the ‘emmeans’ package in R ^100^. In addition to comparing differences in alpha diversity between strains, we compared differences in bacterial community composition by host fungal strain. To do this, we calculated Bray-Curtis dissimilarities between samples using the bacterial community data. For the Bray-Curtis dissimilarity matrix, we explored the effect of host fungal strain identity on bacterial community composition using a permutational analysis of variance (PERMANOVA)(function adonis2 in the ‘vegan’ package in R ^102^).

### Free amino acid quantification

To quantify free amino acids in CCA, we inoculated 150,000 CFUs at a volume of 100 µl per plate containing 20 ml of CCA. We used the same strains as in phenotypic and *in vitro* community assays, as well as 100 µl of PBS as a control. Four replicate plates per strain were used. After incubating for seven days at 24°C in the dark, plates were shipped to Creative Proteomics (Shirley, NY) for free amino acid analysis. A 20 mg subsample was collected for methanol:chloroform extraction. Those samples were run on an HPLC to detect and quantify amino acids. Standard curves were performed with known concentrations of Ala, Arg, Asn, Asp, Cys, Cys2, Gln, Glu, His, HysPro, Ile, Leu, Lys, Met, Orn, Phe, Pro, Ser, Tau, The, Trp, Tyr, Val, and GABA. Concentration of amino acids in the samples were calculated based on the peak of the samples relative to standard curves, with results measured in µ/g. We measured the differences in free amino acid composition by strain by first calculating a Bray-Curtis dissimilarity matrix based on amino acid composition associated with fungal strains. To test the effect of fungal strain identity on amino acid composition, we performed a PERMANOVA using the ‘Vegan’ package in R ^102^ on the distance matrix and visualized the results with a non-metric-multidimensional scaling (NMDS) ordination-based approach.

## REFERENCES

1. Hohl, T.M., Rivera, A., and Pamer, E.G. (2006). Immunity to fungi. Curr. Opin. Immunol. 18, 465–472.

2. Rosa, I., Simoncelli, F., Fagotti, A., and Pascolini, R. (2007). Ecology: The proximate cause of frog declines? Nature 447, E4–E5.

3. Hoyt, J., Kilpatrick, A., and Langwig, K.E. (2021). Ecology and impacts of white-nose syndrome on bats. Nat. Rev. Microbiol. 19, 196–210.

4. Manzoni, M., and Rollini, M. (2002). Biosynthesis and biotechnological production of statins by filamentous fungi and application of these cholesterol-lowering drugs. Appl. Microbiol. Biotechnol. 58, 555–564.

5. Martínez-García, L.B., De Deyn, G.B., Pugnaire, F.I., Kothamasi, D., and van der Heijden, M.G.A. (2017). Symbiotic soil fungi enhance ecosystem resilience to climate change. Glob. Chang. Biol. 23, 5228–5236.

6. Chow, N.A., Muñoz, J.F., Gade, L., Berkow, E.L., Li, X., Welsh, R.M., Forsberg, K., Lockhart, S.R., Adam, R., Alanio, A., et al. (2020). Tracing the evolutionary history and global expansion of Candida auris using population genomic analyses. MBio 11. 10.1128/mBio.03364-19.

7. Ropars, J., Didiot, E., Rodríguez de la Vega, R.C., Bennetot, B., Coton, M., Poirier, E., Coton, E., Snirc, A., Le Prieur, S., and Giraud, T. (2020). Domestication of the emblematic white cheese-making fungus Penicillium camemberti and its diversification into two varieties. Curr. Biol. 30, 4441–4453.e4.

8. Stalder, L., Oggenfuss, U., Mohd-Assaad, N., and Croll, D. (2023). The population genetics of adaptation through copy number variation in a fungal plant pathogen. Mol. Ecol. 32, 2443–2460.

9. McDonald, B.A., and Linde, C. (2002). Pathogen population genetics, evolutionary potential, and durable resistance. Annu. Rev. Phytopathol. 40, 349–379.

10. Cissé, O.H., Ma, L., Wei Huang, D., Khil, P.P., Dekker, J.P., Kutty, G., Bishop, L., Liu, Y., Deng, X., Hauser, P.M., et al. (2018). Comparative population genomics analysis of the mammalian fungal pathogen Pneumocystis. MBio 9. 10.1128/mBio.00381-18.

11. Yuan, Z., Wu, Q., Xu, L., Druzhinina, I.S., Stukenbrock, E.H., Nieuwenhuis, B.P.S., Zhong, Z., Liu, Z.-J., Wang, X., Cai, F., et al. (2022). Genomic landscape of a relict fir-associated fungus reveals rapid convergent adaptation towards endophytism. ISME J. 16, 1294–1305.

12. Sønstebø, J.H., Trucchi, E., Nordén, J., Skrede, I., Miettinen, O., Haridas, S., Pangilinan, J., Grigoriev, I.V., Martin, F., Kauserud, H., et al. (2022). Population genomics of a forest fungus reveals high gene flow and climate adaptation signatures. Mol. Ecol. 31, 1963–1979.

13. Plissonneau, C., Benevenuto, J., Mohd-Assaad, N., Fouché, S., Hartmann, F.E., and Croll, D. (2017). Using population and comparative genomics to understand the genetic basis of effector-driven fungal pathogen evolution. Front. Plant Sci. 8, 119.

14. Tso, G.H.W., Reales-Calderon, J.A., Tan, A.S.M., Sem, X., Le, G.T.T., Tan, T.G., Lai, G.C., Srinivasan, K.G., Yurieva, M., Liao, W., et al. (2018). Experimental evolution of a fungal pathogen into a gut symbiont. Science 362, 589–595.

15. Schoustra, S.E., Debets, A.J.M., Slakhorst, M., and Hoekstra, R.F. (2006). Reducing the cost of resistance; experimental evolution in the filamentous fungus Aspergillus nidulans. J. Evol. Biol. 19, 1115–1127.

16. Fisher, K.J., and Lang, G.I. (2016). Experimental evolution in fungi: An untapped resource. Fungal Genet. Biol. 94, 88–94.

17. Zeyl, C. (2006). Experimental evolution with yeast. FEMS Yeast Res. 6, 685–691.

18. Kawecki, T.J., Lenski, R.E., Ebert, D., Hollis, B., Olivieri, I., and Whitlock, M.C. (2012). Experimental evolution. Trends Ecol. Evol. 27, 547–560.

19. Martín, J., and Coton, M. (2017). Blue cheese: Microbiota and fungal metabolites. Fermented foods in health and disease prevention, 275–303.

20. Özer, B. (2014). Cheese Microbiology. In Dairy Microbiology and Biochemistry (CRC Press), pp. 127–147.

21. Kable, M.E., Srisengfa, Y., Laird, M., Zaragoza, J., McLeod, J., Heidenreich, J., and Marco, M.L. (2016). The core and seasonal Microbiota of raw bovine milk in tanker trucks and the impact of transfer to a milk processing facility. MBio 7. 10.1128/mBio.00836-16.

22. Bokulich, N.A., and Mills, D.A. (2013). Facility-specific “house” microbiome drives microbial landscapes of artisan cheesemaking plants. Appl. Environ. Microbiol. 79, 5214–5223.

23. Sun, L., and D’Amico, D.J. (2021). Composition, succession, and source tracking of microbial communities throughout the traditional production of a farmstead cheese. mSystems 6, e0083021.

24. Dumas, E., Feurtey, A., Rodríguez de la Vega, R.C., Le Prieur, S., Snirc, A., Coton, M., Thierry, A., Coton, E., Le Piver, M., Roueyre, D., et al. (2020). Independent domestication events in the blue-cheese fungus Penicillium roqueforti. Mol. Ecol. 29, 2639–2660.

25. Wolfe, B.E., Button, J.E., Santarelli, M., and Dutton, R.J. (2014). Cheese rind communities provide tractable systems for in situ and in vitro studies of microbial diversity. Cell 158, 422–433.

26. Biango-Daniels, M.N., and Wolfe, B.E. (2021). American artisan cheese quality and spoilage: A survey of cheesemakers’ concerns and needs. J. Dairy Sci. 104, 6283–6294.

27. Marcellino, N., and Benson, D.R. (2013). The Good, the Bad, and the Ugly: Tales of Mold-Ripened Cheese. Microbiol Spectr 1. 10.1128/microbiolspec.CM-0005-12.

28. Bodinaku, I., Shaffer, J., Connors, A.B., Steenwyk, J.L., Biango-Daniels, M.N., Kastman, E.K., Rokas, A., Robbat, A., and Wolfe, B.E. (2019). Rapid phenotypic and metabolomic domestication of wild Penicillium molds on cheese. MBio 10. 10.1128/mBio.02445-19.

29. Ropars, J., Cruaud, C., Lacoste, S., and Dupont, J. (2012). A taxonomic and ecological overview of cheese fungi. Int. J. Food Microbiol. 155, 199–210.

30. Gonçalves, V.N., Campos, L.S., Melo, I.S., Pellizari, V.H., Rosa, C.A., and Rosa, L.H. (2013). Penicillium solitum: a mesophilic, psychrotolerant fungus present in marine sediments from Antarctica. Polar Biol. 36, 1823–1831.

31. Visagie, C.M., Magistà, D., Ferrara, M., Balocchi, F., Duong, T.A., Eichmeier, A., Gramaje, D., Aylward, J., Baker, S.E., Barnes, I., et al. (2023). IMA genome-F18 : The re-identification of Penicillium genomes available in NCBI and draft genomes for Penicillium species from dry cured meat, Penicillium biforme, P. brevicompactum, P. solitum, and P. cvjetkovicii, Pewenomyces kutranfy, Pew. lalenivora, Pew. tapulicola, Pew. kalosus, Teratosphaeria carnegiei, and Trichoderma atroviride SC1: The re-identification of Penicillium genomes available in NCBI and draft genomes for Penicillium species from dry cured meat, Penicillium biforme, P. brevicompactum, P. solitum, and P. cvjetkovicii, Pewenomyces kutranfy, Pew. lalenivora, Pew. tapulicola, Pew. kalosus, Teratosphaeria carnegiei, and Trichoderma atroviride SC1. IMA Fungus 14, 21.

32. Kastman, E.K., Kamelamela, N., Norville, J.W., Cosetta, C.M., Dutton, R.J., and Wolfe, B.E. (2016). Biotic interactions shape the ecological distributions of staphylococcus species. MBio 7, e01157–16.

33. Pierce, E.C., Morin, M., Little, J.C., Liu, R., Tannous, J., Keller, N., Pogliano, K., Wolfe, B., Sanchez, L.M., and Dutton, R. (2020). Bacterial-fungal interactions revealed by genome-wide analysis of bacterial mutant fitness. Nat. Microbiol. 6, 87–102.

34. Butler, M.J., Day, A.W., Henson, J.M., and Money, N.P. (2001). Pathogenic properties of fungal melanins. Mycologia 93, 1–8.

35. Eisenman, H.C., and Casadevall, A. (2012). Synthesis and assembly of fungal melanin. Appl. Microbiol. Biotechnol. 93, 931–940.

36. Cleere, M.M., Novodvorska, M., Geib, E., Whittaker, J., Dalton, H., Salih, N., Hewitt, S., Kokolski, M., Brock, M., and Dyer, P.S. (2024). New colours for old in the blue-cheese fungus Penicillium roqueforti. Npj Sci. Food 8, 3.

37. Perez-Cuesta, U., Aparicio-Fernandez, L., Guruceaga, X., Martin-Souto, L., Abad-Diaz-de-Cerio, A., Antoran, A., Buldain, I., Hernando, F.L., Ramirez-Garcia, A., and Rementeria, A. (2020). Melanin and pyomelanin in Aspergillus fumigatus: from its genetics to host interaction. Int. Microbiol. 23, 55–63.

38. Jia, S.-L., Chi, Z., Chen, L., Liu, G.-L., Hu, Z., and Chi, Z.-M. (2021). Molecular evolution and regulation of DHN melanin-related gene clusters are closely related to adaptation of different melanin-producing fungi. Genomics 113, 1962–1975.

39. Muszewska, A., Steczkiewicz, K., Stepniewska-Dziubinska, M., and Ginalski, K. (2019). Transposable elements contribute to fungal genes and impact fungal lifestyle. Sci. Rep. 9, 4307.

40. Muszewska, A., Hoffman-Sommer, M., and Grynberg, M. (2011). LTR retrotransposons in fungi. PLoS One 6, e29425.

41. Cai, X., Lin, R., Liang, J., King, G.J., Wu, J., and Wang, X. (2022). Transposable element insertion: a hidden major source of domesticated phenotypic variation in Brassica rapa. Plant Biotechnol. J. 20, 1298–1310.

42. Britten, R.J. (2010). Transposable element insertions have strongly affected human evolution. Proc. Natl. Acad. Sci. U. S. A. 107, 19945–19948.

43. Oggenfuss, U., Badet, T., Wicker, T., Hartmann, F.E., Singh, N.K., Abraham, L., Karisto, P., Vonlanthen, T., Mundt, C., McDonald, B.A., et al. (2021). A population-level invasion by transposable elements triggers genome expansion in a fungal pathogen. Elife 10. 10.7554/eLife.69249.

44. Krishnan, P., Meile, L., Plissonneau, C., Ma, X., Hartmann, F.E., Croll, D., McDonald, B.A., and Sánchez-Vallet, A. (2018). Transposable element insertions shape gene regulation and melanin production in a fungal pathogen of wheat. BMC Biol. 16, 78.

45. Cordero, R.J., and Casadevall, A. (2017). Functions of fungal melanin beyond virulence. Fungal Biol. Rev. 31, 99–112.

46. Blachowicz, A., Raffa, N., Bok, J.W., Choera, T., Knox, B., Lim, F.Y., Huttenlocher, A., Wang, C.C.C., Venkateswaran, K., and Keller, N.P. (2020). Contributions of spore secondary metabolites to UV-C protection and virulence vary in different Aspergillus fumigatus strains. MBio 11. 10.1128/mBio.03415-19.

47. Bilandžija, H., Laslo, M., Porter, M., and Fong, D. (2017). Melanization in response to wounding is ancestral in arthropods and conserved in albino cave species. Sci. Rep. 7. 10.1038/s41598-017-17471-2.

48. Bilandžija, H., Cetković, H., and Jeffery, W.R. (2012). Evolution of albinism in cave planthoppers by a convergent defect in the first step of melanin biosynthesis. Evol. Dev. 14, 196–203.

49. Tannous, J., Cosetta, C.M., Drott, M.T., Rush, T.A., Abraham, P.E., Giannone, R.J., Keller, N.P., and Wolfe, B.E. (2023). LaeA-regulated fungal traits mediate bacterial community assembly. MBio 14, e0076923.

50. Cosetta, C.M., Kfoury, N., Robbat, A., and Wolfe, B.E. (2020). Fungal volatiles mediate cheese rind microbiome assembly. Environ. Microbiol. 22, 4745–4760.

51. Zhang, Y., Kastman, E., Guasto, J., and Wolfe, B. (2018). Fungal networks shape dynamics of bacterial dispersal and community assembly in cheese rind microbiomes. Nat. Commun. 9. 10.1038/s41467-017-02522-z.

52. Monnet, C., Landaud, S., Bonnarme, P., and Swennen, D. (2015). Growth and adaptation of microorganisms on the cheese surface. FEMS Microbiol. Lett. 362, 1–9.

53. Fox, P. (2014). Cheese: Chemistry, physics and microbiology: General aspects 3rd ed. P. F. Fox, P. L. H. McSweeney, T. M. Cogan, and T. P. Guinee, eds. (Academic Press).

54. Marsit, S., Leducq, J.-B., Durand, É., Marchant, A., Filteau, M., and Landry, C.R. (2017). Evolutionary biology through the lens of budding yeast comparative genomics. Nat. Rev. Genet. 18, 581–598.

55. Gibbons, J.G., Salichos, L., Slot, J.C., Rinker, D.C., McGary, K.L., King, J.G., Klich, M.A., Tabb, D.L., McDonald, W.H., and Rokas, A. (2012). The evolutionary imprint of domestication on genome variation and function of the filamentous fungus Aspergillus oryzae. Curr. Biol. 22, 1403–1409.

56. Machida, M., Yamada, O., and Gomi, K. (2008). Genomics of Aspergillus oryzae: learning from the history of Koji mold and exploration of its future. DNA Res. 15, 173–183.

57. Abe, K., Gomi, K., Hasegawa, F., and Machida, M. (2006). Impact of Aspergillus oryzae genomics on industrial production of metabolites. Mycopathologia 162, 143–153.

58. Singaravelan, N., Grishkan, I., Beharav, A., Wakamatsu, K., Ito, S., and Nevo, E. (2008). Adaptive melanin response of the soil fungus Aspergillus niger to UV radiation stress at “Evolution Canyon”, Mount Carmel, Israel. PLoS One 3, e2993.

59. Pereira, D., Croll, D., Brunner, P.C., and McDonald, B.A. (2020). Natural selection drives population divergence for local adaptation in a wheat pathogen. Fungal Genet. Biol. 141, 103398.

60. Desjardins, C.A., Giamberardino, C., Sykes, S.M., Yu, C.-H., Tenor, J.L., Chen, Y., Yang, T., Jones, A.M., Sun, S., Haverkamp, M.R., et al. (2017). Population genomics and the evolution of virulence in the fungal pathogen Cryptococcus neoformans. Genome Res. 27, 1207–1219.

61. Robertson, K.L., Mostaghim, A., Cuomo, C.A., Soto, C.M., Lebedev, N., Bailey, R.F., and Wang, Z. (2012). Adaptation of the black yeast Wangiella dermatitidis to ionizing radiation: molecular and cellular mechanisms. PLoS One 7, e48674.

62. Yuan, Z., Druzhinina, I.S., Gibbons, J.G., Zhong, Z., Van de Peer, Y., Rodriguez, R.J., Liu, Z., Wang, X., Wei, H., Wu, Q., et al. (2021). Divergence of a genomic island leads to the evolution of melanization in a halophyte root fungus. ISME J. 15, 3468–3479.

63. Protas, M., Conrad, M., Gross, J.B., Tabin, C., and Borowsky, R. (2007). Regressive evolution in the Mexican cave tetra, Astyanax mexicanus. Curr. Biol. 17, 452–454.

64. Liu, W., Golovatch, S., Wesener, T., and Tian, M. (2017). Convergent evolution of unique morphological adaptations to a subterranean environment in cave millipedes (Diplopoda). PLoS One 12, e0170717.

65. Soares, D., and Niemiller, M.L. (2020). Extreme adaptation in caves. Anat. Rec. (Hoboken) 303, 15–23.

66. Fouché, S., Oggenfuss, U., Chanclud, E., and Croll, D. (2022). A devil’s bargain with transposable elements in plant pathogens. Trends Genet. 38, 222–230.

67. Lebreton, A., Tang, N., Kuo, A., LaButti, K., Andreopoulos, W., Drula, E., Miyauchi, S., Barry, K., Clum, A., Lipzen, A., et al. (2022). Comparative genomics reveals a dynamic genome evolution in the ectomycorrhizal milk-cap (Lactarius) mushrooms. New Phytol. 235, 306–319.

68. Torres, D.E., Thomma, B.P.H.J., and Seidl, M.F. (2021). Transposable elements contribute to genome dynamics and gene expression variation in the fungal plant pathogen Verticillium dahliae. Genome Biol. Evol. 13. 10.1093/gbe/evab135.

69. Dallaire, A., Manley, B.F., Wilkens, M., Bista, I., Quan, C., Evangelisti, E., Bradshaw, C.R., Ramakrishna, N.B., Schornack, S., Butter, F., et al. (2021). Transcriptional activity and epigenetic regulation of transposable elements in the symbiotic fungus Rhizophagus irregularis. Genome Res. 31, 2290–2302.

70. Gluck-Thaler, E., Ralston, T., Konkel, Z., Ocampos, C.G., Ganeshan, V.D., Dorrance, A.E., Niblack, T.L., Wood, C.W., Slot, J.C., Lopez-Nicora, H.D., et al. (2022). Giant Starship elements mobilize accessory genes in fungal genomes. Mol. Biol. Evol. 39. 10.1093/molbev/msac109.

71. Bourque, G., Burns, K.H., Gehring, M., Gorbunova, V., Seluanov, A., Hammell, M., Imbeault, M., Izsvák, Z., Levin, H.L., Macfarlan, T.S., et al. (2018). Ten things you should know about transposable elements. Genome Biol. 19, 199.

72. Zhang, X., Meng, L., Liu, B., Hu, Y., Cheng, F., Liang, J., Aarts, M.G.M., Wang, X., and Wu, J. (2015). A transposon insertion in FLOWERING LOCUS T is associated with delayed flowering in Brassica rapa. Plant Sci. 241, 211–220.

73. O’Donnell, S., Rezende, G., Vernadet, J.-P., Snirc, A., and Ropars, J. (2024). Harbouring Starships: the accumulation of large horizontal gene transfers in domesticated and pathogenic fungi. bioRxiv, 2024.07. 03.601904. 10.1101/2024.07.03.601904.

74. Hedges, D.J., and Deininger, P.L. (2007). Inviting instability: Transposable elements, double-strand breaks, and the maintenance of genome integrity. Mutat. Res. 616, 46–59.

75. Ropars, J., and Giraud, T. (2022). Convergence in domesticated fungi used for cheese and dry-cured meat maturation: beneficial traits, genomic mechanisms, and degeneration. Curr. Opin. Microbiol. 70, 102236.

76. Ropars, J., Caron, T., Lo, Y.-C., Bennetot, B., and Giraud, T. (2020). The domestication of Penicillium cheese fungi. C. R. Biol. 343, 155–176.

77. Arrigan, D., Kothe, C.I., Oliverio, A., Evans, J.D., and Wolfe, B.E. (2024). Novel fermentations integrate traditional practice and rational design of fermented-food microbiomes. Curr. Biol. 34, R1094–R1108.

78. Gibbons, J.G., and Rinker, D.C. (2015). The genomics of microbial domestication in the fermented food environment. Curr. Opin. Genet. Dev. 35, 1–8.

79. Lo, Y.-C., Bruxaux, J., Rodríguez de la Vega, R.C., O’Donnell, S., Snirc, A., Coton, M., Le Piver, M., Le Prieur, S., Roueyre, D., Dupont, J., et al. (2023). Domestication in dry-cured meat Penicillium fungi: Convergent specific phenotypes and horizontal gene transfers without strong genetic subdivision. Evol. Appl. 16, 1637–1660.

80. Cosetta, C.M., and Wolfe, B.E. (2020). Deconstructing and Reconstructing Cheese Rind Microbiomes for Experiments in Microbial Ecology and Evolution. Curr. Protoc. Microbiol. 56, e95.

81. Wu, H.-M. (2022). QIAGEN® DNeasy® PowerSoil® Pro v1. 10.17504/protocols.io.bp2l69411lqe/v1.

82. Menzel, P., Ng, K.L., and Krogh, A. (2016). Fast and sensitive taxonomic classification for metagenomics with Kaiju. Nat. Commun. 7, 11257.

83. Arkin, A.P., Cottingham, R.W., Henry, C.S., Harris, N.L., Stevens, R.L., Maslov, S., Dehal, P., Ware, D., Perez, F., Canon, S., et al. (2018). KBase: The United States department of energy systems biology knowledgebase. Nat. Biotechnol. 36, 566–569.

84. Koren, S., Walenz, B.P., Berlin, K., Miller, J.R., Bergman, N.H., and Phillippy, A.M. (2017). Canu: scalable and accurate long-read assembly via adaptive k-mer weighting and repeat separation. Genome Res. 27, 722–736.

85. Walker, B.J., Abeel, T., Shea, T., Priest, M., Abouelliel, A., Sakthikumar, S., Cuomo, C.A., Zeng, Q., Wortman, J., Young, S.K., et al. (2014). Pilon: an integrated tool for comprehensive microbial variant detection and genome assembly improvement. PLoS One 9, e112963.

86. Li, H., and Durbin, R. (2010). Fast and accurate long-read alignment with Burrows-Wheeler transform. Bioinformatics 26, 589–595.

87. Li, H., Handsaker, B., Wysoker, A., Fennell, T., Ruan, J., Homer, N., Marth, G., Abecasis, G., Durbin, R., and 1000 Genome Project Data Processing Subgroup (2009). The Sequence Alignment/Map format and SAMtools. Bioinformatics 25, 2078–2079.

88. Humann, J.L., Lee, T., Ficklin, S., and Main, D. (2019). Structural and Functional Annotation of Eukaryotic Genomes with GenSAS. Methods Mol Biol 1962, 29–51.

89. Stanke, M., and Waack, S. (2003). Gene prediction with a hidden Markov model and a new intron submodel. Bioinformatics 19 Suppl 2, ii215–ii225.

90. Gauch, H.G., Jr, Qian, S., Piepho, H.-P., Zhou, L., and Chen, R. (2019). Consequences of PCA graphs, SNP codings, and PCA variants for elucidating population structure. PLoS One 14, e0218306.

91. Edgar, R.C. (2004). MUSCLE: multiple sequence alignment with high accuracy and high throughput. Nucleic Acids Res. 32, 1792–1797.

92. Flynn, J.M., Hubley, R., Goubert, C., Rosen, J., Clark, A.G., Feschotte, C., and Smit, A.F. (2020). RepeatModeler2 for automated genomic discovery of transposable element families. Proc. Natl. Acad. Sci. U. S. A. 117, 9451–9457.

93. Love, M.I., Huber, W., and Anders, S. (2014). Moderated estimation of fold change and dispersion for RNA-seq data with DESeq2. Genome Biol. 15, 550.

94. Xie, C., Mao, X., Huang, J., Ding, Y., Wu, J., Dong, S., Kong, L., Gao, G., Li, C.-Y., and Wei, L. (2011). KOBAS 2.0: a web server for annotation and identification of enriched pathways and diseases. Nucleic Acids Res. 39, W316–W322.

95. ggplot2 https://ggplot2.tidyverse.org.

96. Hartmann, T., Dümig, M., Jaber, B.M., Szewczyk, E., Olbermann, P., Morschhäuser, J., and Krappmann, S. (2010). Validation of a self-excising marker in the human pathogen Aspergillus fumigatus by employing the beta-rec/six site-specific recombination system. Appl. Environ. Microbiol. 76, 6313–6317.

97. Nickles, G., Ludwikoski, I., Bok, J.W., and Keller, N.P. (2021). Comprehensive guide to extracting and expressing fungal secondary metabolites with Aspergillus fumigatus as a case study. Curr. Protoc. 1, e321.

98. Fernandez, C.W., and Koide, R.T. (2014). Initial melanin and nitrogen concentrations control the decomposition of ectomycorrhizal fungal litter. Soil Biol. Biochem. 77, 150–157.

99. Siletti, C.E., Zeiner, C.A., and Bhatnagar, J.M. (2017). Distributions of fungal melanin across species and soils. Soil Biol. Biochem. 113, 285–293.

100. Lenth, {russell V., Banfai, B., Bolker, B., Buerkner, P., Giné-Vázquez, I., Herve, M., Jung, M., Love, J., Miguez, F., Piaskowski, J., et al. (2012). Estimated Marginal Means, aka Least-Squares Means. CRAN.

101. Fox, J., and Weisberg, S. (2018). An R companion to applied regression 3rd ed. (SAGE Publications).

102. Dixon, P. (2003). VEGAN, a package of R functions for community ecology. J. Veg. Sci. 14, 927–930.

