## Supplementary material for "Transposable elements facilitate the unintentional domestication of a cheese-associated *Penicillium* mold": Figure S

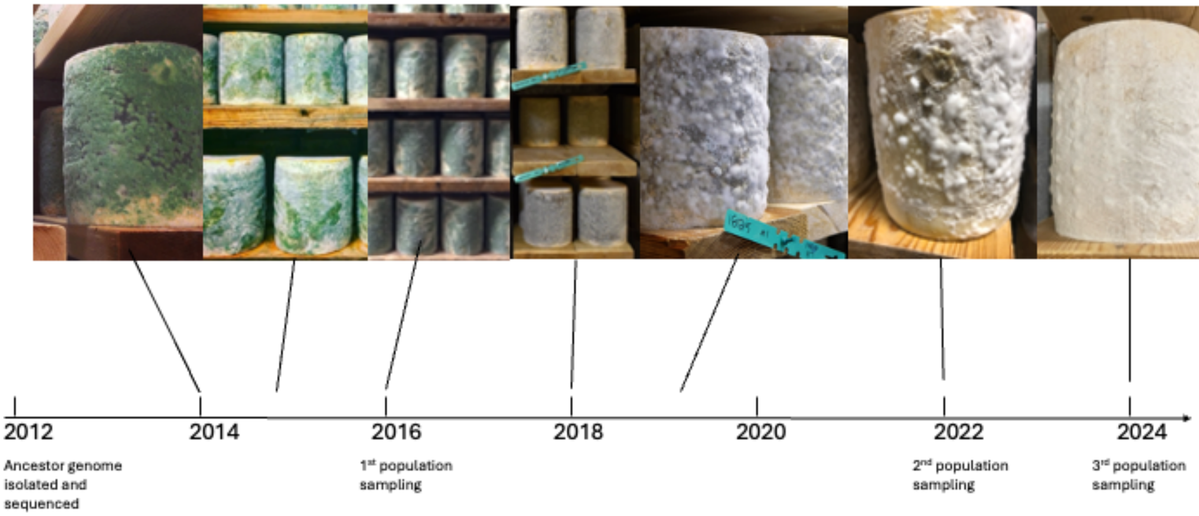


**Figure S1:** Photographic documentation of the green to white phenotypic shift of Bayley Hazen Blue cheese and associated *Penicillium solitum*. All photographs were taken within the same vault of the same cheese cave. Photographs from 2014 were taken by Bedford Cheese Shop (used with permission), from 2015 and 2018 by Jasper Hill Farms (used with permission), and 2019 by Meg Maker (used with permission). The initially bright green color of *P. solitum* can fade to a grey/brown color (observed in 2019 and 2022 photos) more characteristic of other fungal melanins. Related to Figure 1.

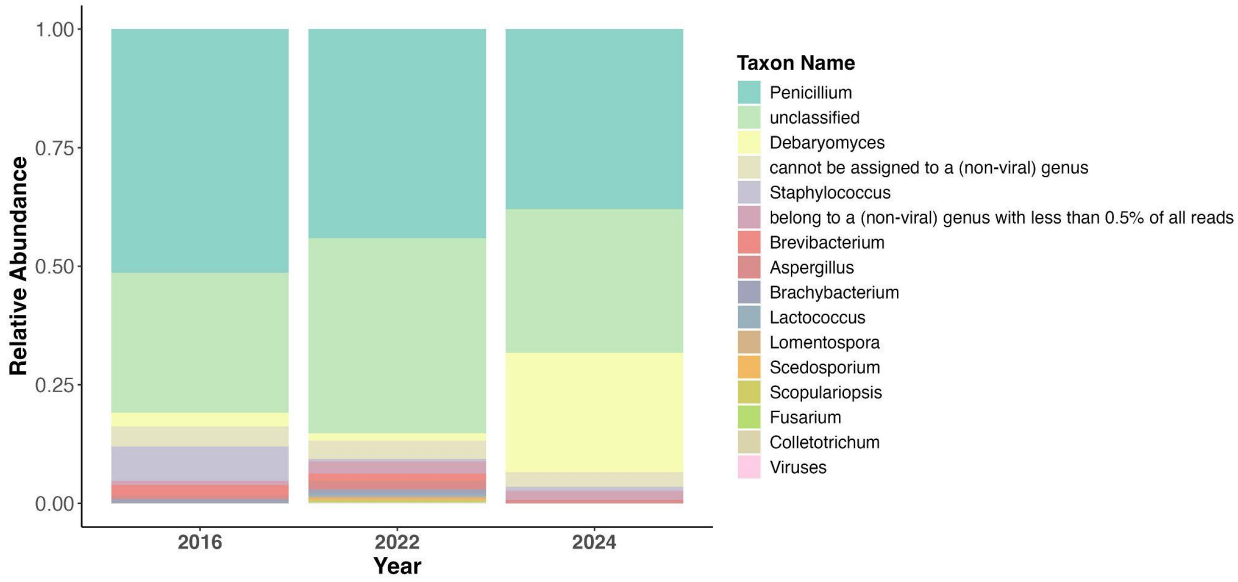


**Figure S2:** Proportion of reads classified to genus level (using kaiju) across the 9 cheese rind samples selected for metagenomic sequencing. Related to Figure 1.

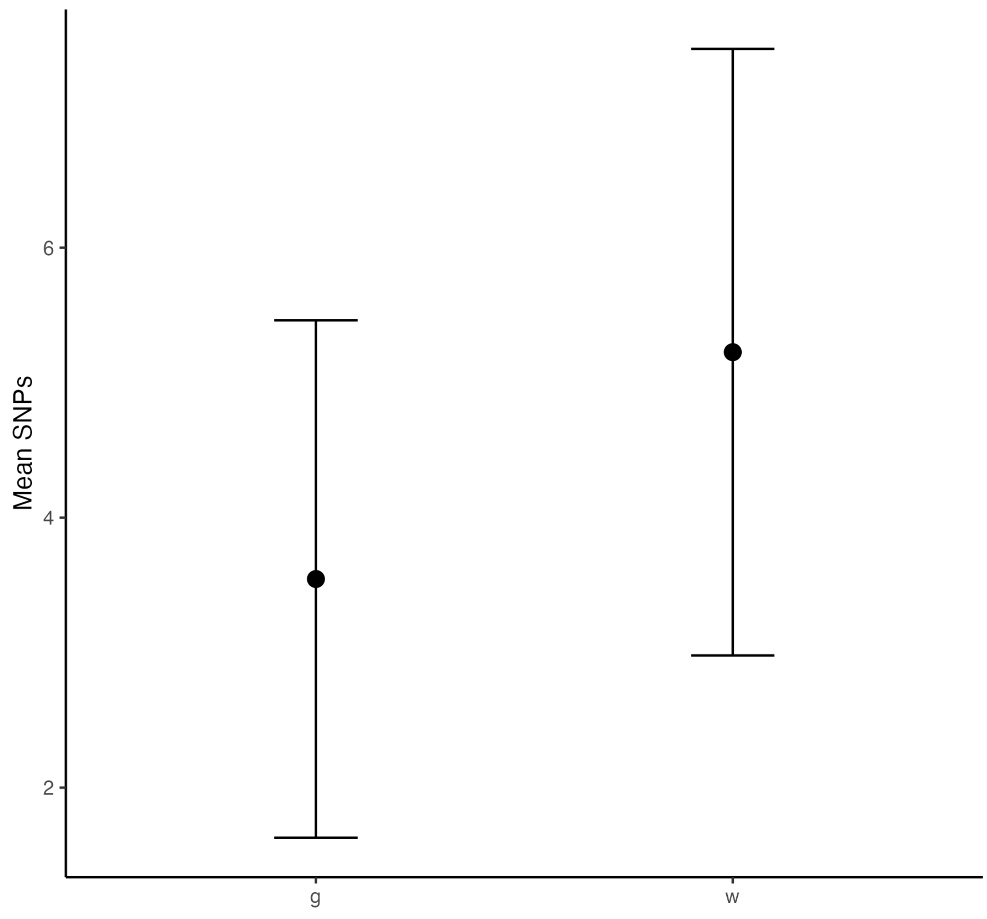


**Figure S3:** Mean number of non-synonymous mutations (SNPs and short Indels) within coding sequences of green (g) and white (w) strains after applying filters of minimum coverage of 20X and variant frequency of 90%. Error bars represent standard error. Related to Figure 2.

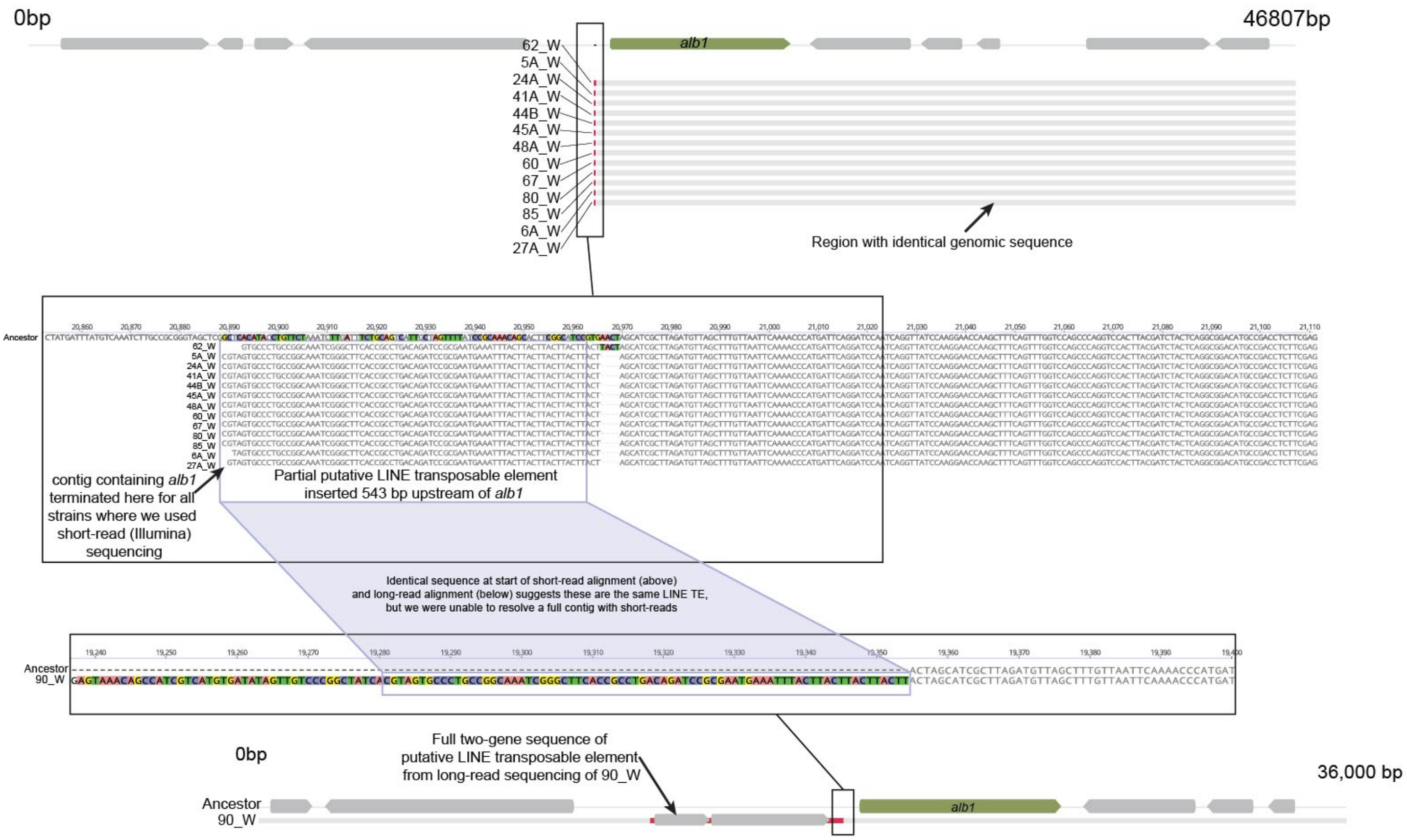


**Figure S4:** Alignments of the *alb1* gene region illustrating putative transposable element insertions in the upstream region of *alb1*. In “zoomed out” alignments, regions in grey are conserved across mutant and ancestor sequences. Areas in red indicate mutations or insertions. The top alignment shows various strains that had a partial insertion of a putative LINE transposable element (TE). Short-read sequencing may not have been able to fully assemble this region because transposable elements are repetitive. When we used a combination of long-read (Nanopore) and short-read (Illumina) sequencing for one of these mutants (90_W; shown in the bottom alignment) we were able to capture the full putative LINE TE, including the two genes encoded by the TE. The area shaded in blue shows an identical sequence between the short-read genomes and hybrid genome for 90_W, suggesting these are the same TE insertion. Related to Figure 2.

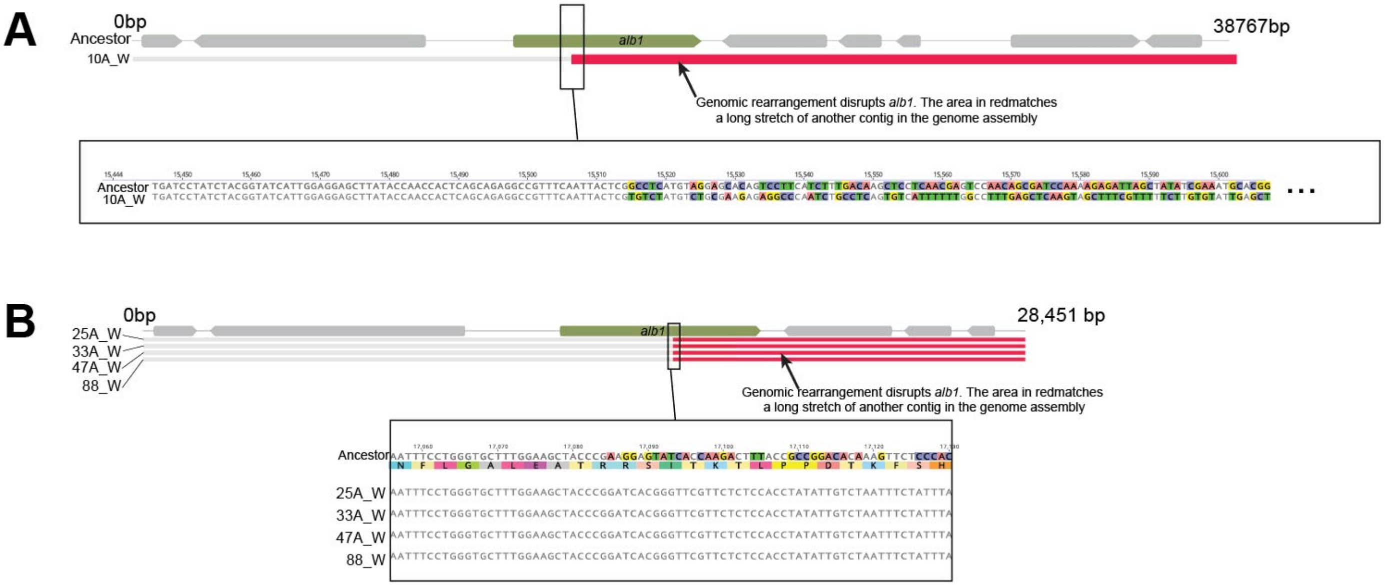


**Figure S5:** Alignments of the *alb1* gene region to illustrate putative genomic rearrangements that disrupted the *alb1* gene. In "zoomed out" alignments, regions in grey are conserved across mutant and ancestor sequences. Areas in red indicate rearrangements where large stretches match other contigs in the genome assembly. Related to Figure 2.

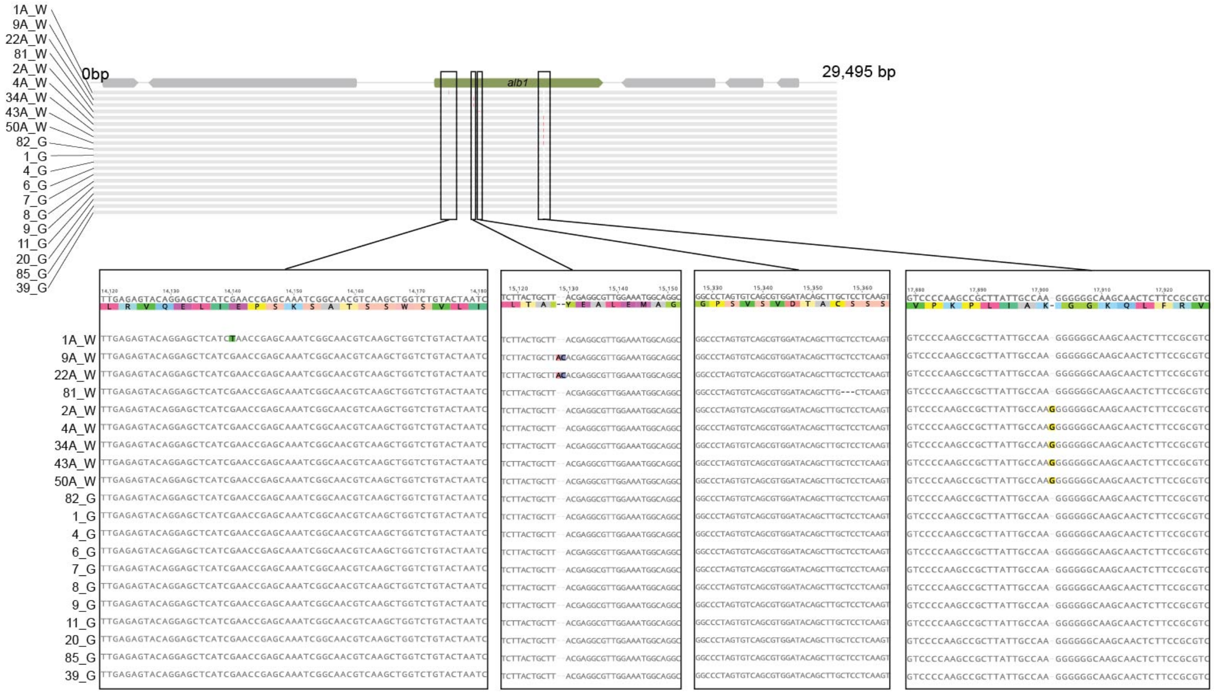


**Figure S6:** Alignments of the *alb1* gene region to illustrate SNPs and indels in various white *P. solitum* strains (with "_W"). Green isolates (with "_G") are also shown in the alignment to illustrate the lack of mutations in these strains. In "zoomed out" alignment, regions in grey are conserved across mutant and ancestor sequences. Areas in red indicate mutations. Related to Figure 2.

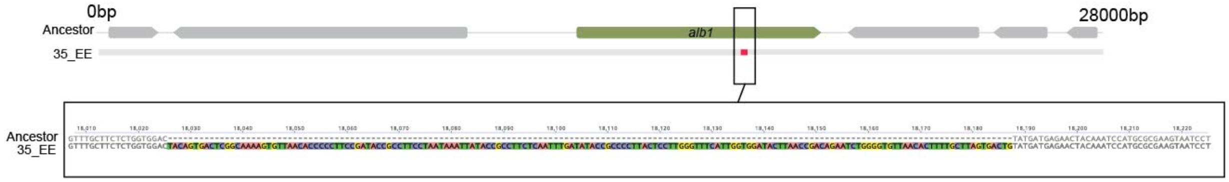


**Figure S7:** Alignment of the *alb1* gene region to highlight a large insertion in the experimentally evolved isolate 35_EE. In "zoomed out" alignments, regions in grey are conserved across mutant and ancestor sequences. The area in red indicates a 162 bp insertion in the alb1 gene. This sequence is repeated in intergenic regions throughout the genome and may be a mobile genetic element. Related to Figure 2.

**A**

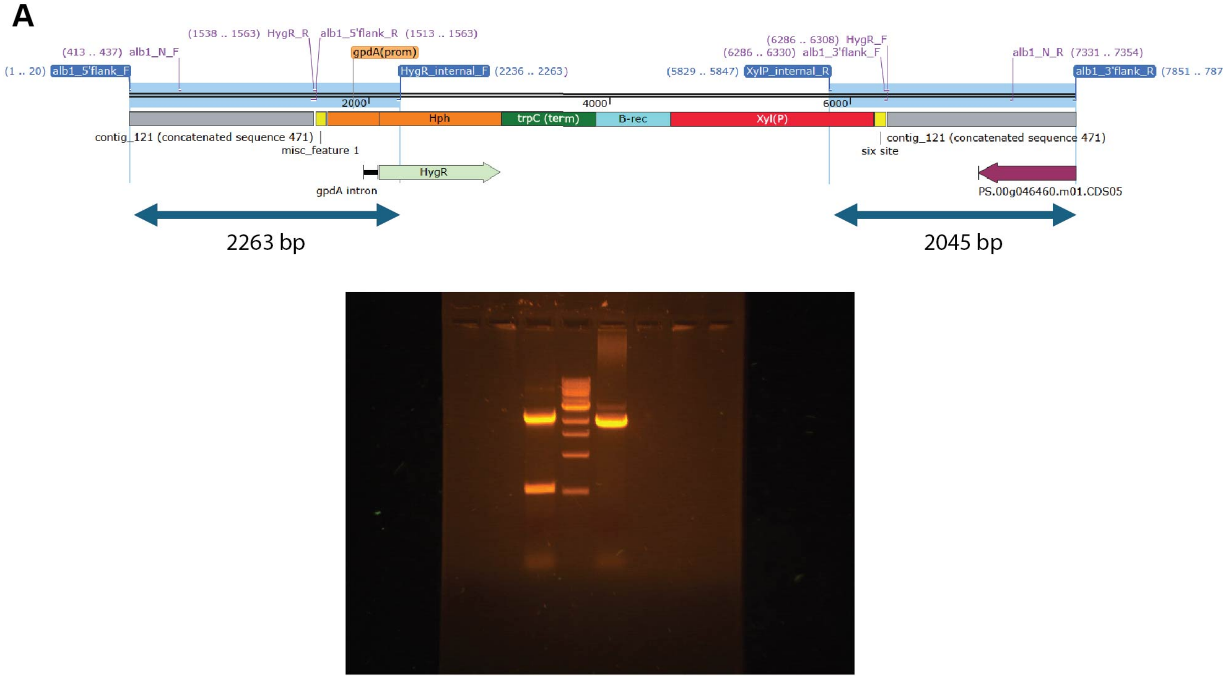


**B**

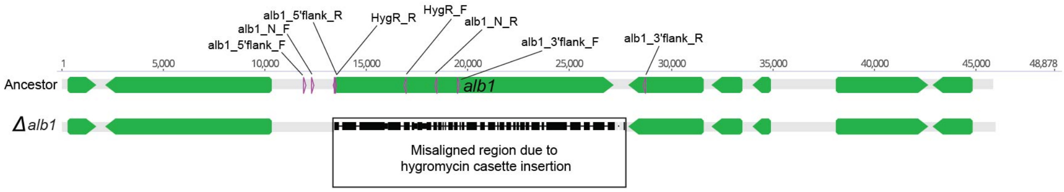


**Figure S8:** Construction and confirmation of an *alb1* knockout in *Penicillium solitum* strain #12. (A) Overview of primers used for constructing the knockout and PCR results to confirm successful deletion. White transformants were screened for *alb1* deletions using two primer pairs, with one primer for each pair in the hygromycin-resistance cassette and the other within the genome outside of the recombined region. One transformant successfully provided both of these amplicons (lane to the left of the ladder). (B) Whole-genome sequencing was used to confirm proper insertion of the hygromycin cassette in the *alb1* region. The alignment shows the Ancestor genome on top and the alb1 knockout on the bottom with a missing *alb1* gene. Related to Figure 2.

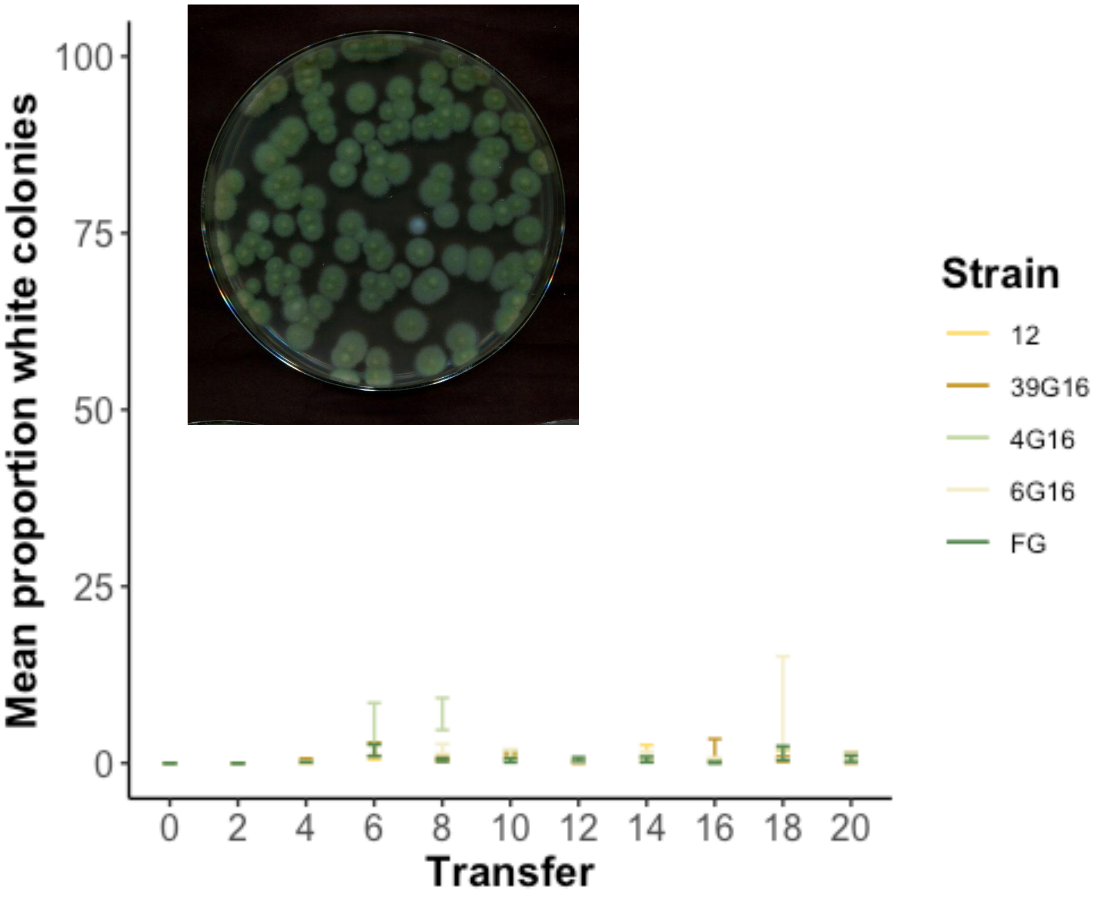


**Figure S9:** White phenotypes emerge after 6 weeks of serial transfers, but stay at a low frequency. Mean proportion of white colonies calculated across 8 biological replicates within five unique green strains of *Penicillium solitum*. Error bars indicate one standard error of the mean. Scanned image is an example of a white colony that emerged in a biological replicate at week 6. Related to Figure 2.

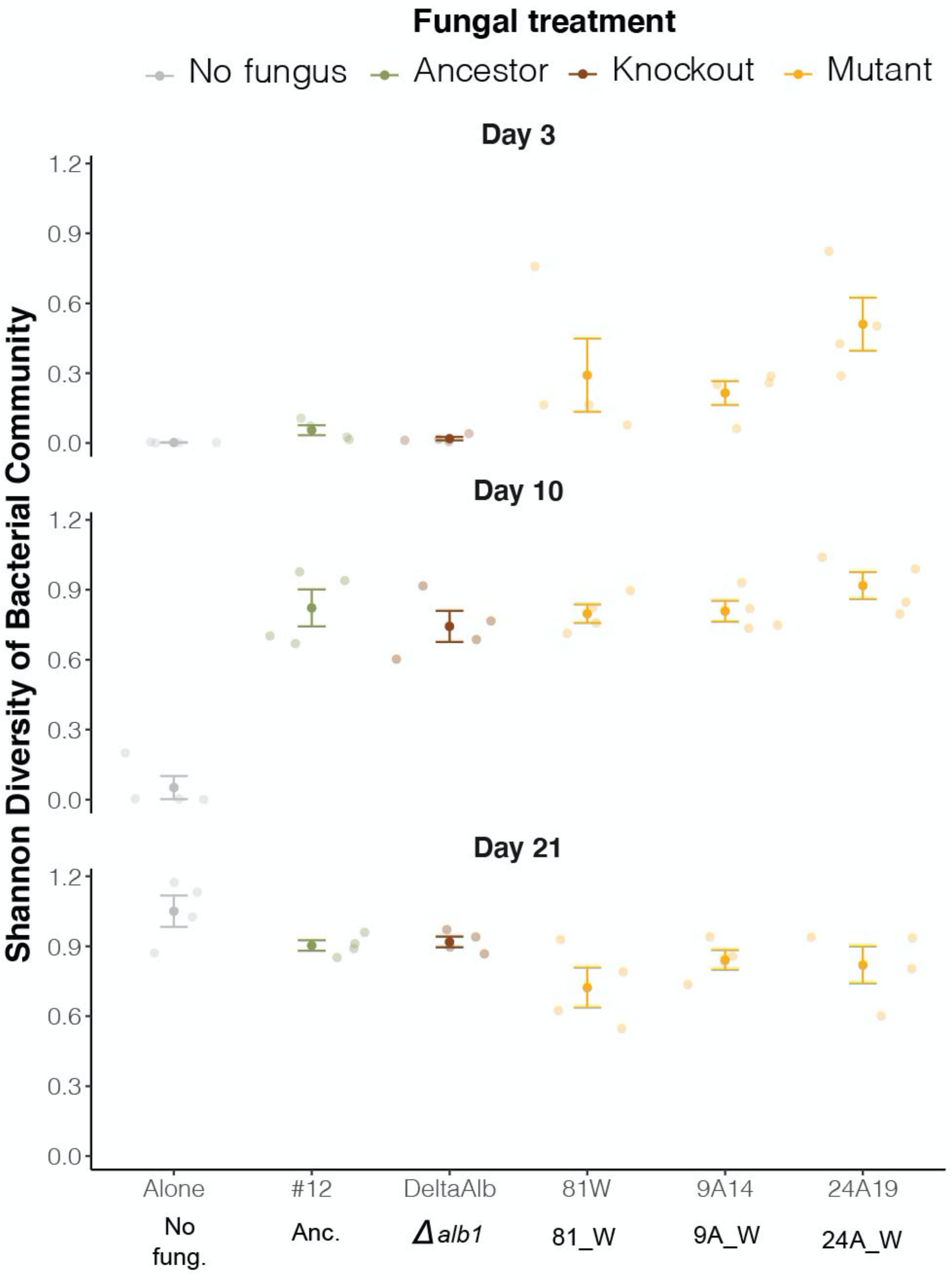


**Figure S10:** Shannon diversity of bacterial communities with different *Penicillium solitum* strains. Related to Figure 4.

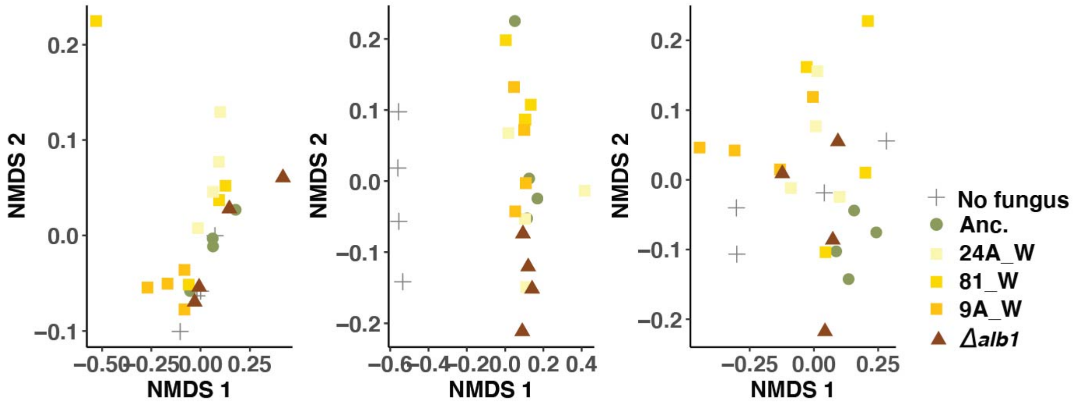


**Figure S11:** Non-metric multidimensional scaling (NMDS) plots of the four-member bacterial communities. Panels are ordered from harvest day 3 (left), 10 (middle) to 21 (right). Related to Figure 4.

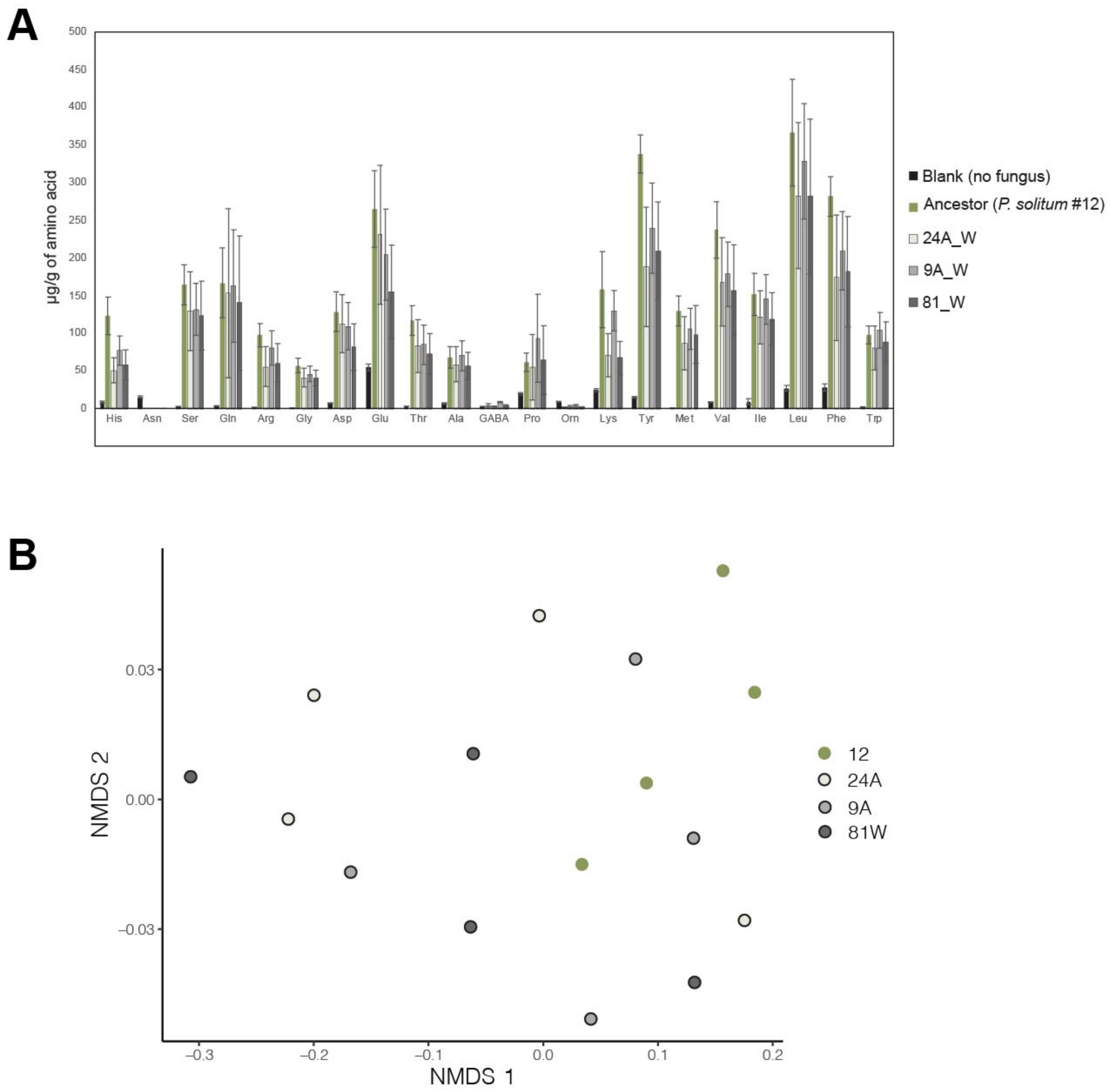


**Figure S12: (A)** Absolute abundance of available free amino acids detected in different strain treatments. Each bar represents mean concentration of amino acids measured. Each error bar represents one standard deviation. **(B)** Non-metric multidimensional scaling (NMDS) plots of available free amino acid composition by different fungal strains. Related to Figure 4.
